# Tfap2b specifies an embryonic melanocyte stem cell that retains adult multi-fate potential

**DOI:** 10.1101/2021.06.18.448859

**Authors:** Alessandro Brombin, Daniel J. Simpson, Jana Travnickova, Hannah Brunsdon, Zhiqiang Zeng, Yuting Lu, Adelaide I.J. Young, Tamir Chandra, E. Elizabeth Patton

**Affiliations:** MRC Human Genetics Unit, Institute of Genetics and Cancer, University of Edinburgh, Western General Hospital Campus, EH4 2XU, Edinburgh, United Kingdom.; CRUK Edinburgh Centre, Institute of Genetics and Cancer, University of Edinburgh, Western General Hospital Campus, EH4 2XU, Edinburgh, United Kingdom.

**Keywords:** Melanocyte stem cells, pigment stem cells, neural crest cells, *tfap2b*, ErbB signaling, melanocyte lineage, scRNA-seq, fate mapping, lineage tracing, chemical genetics, zebrafish

## Abstract

Melanocytes, our pigment producing cells, are replenished from multiple stem cell niches in adult tissues. Although pigmentation traits are known risk-factors for melanoma, we know little about melanocyte stem cell (McSC) populations other than hair follicle McSCs, and lack key lineage markers with which to identify McSCs and study their function. Here, we discover that Tfap2b, and a select set of its target genes, specifies an McSC population at the dorsal root ganglia in zebrafish. Functionally, Tfap2b is required for only a few late-stage embryonic melanocytes, and instead is essential for McSC-dependent melanocyte regeneration. Fate-mapping data reveal that *tfap2b*-expressing McSCs have multi-fate potential, and are the cell-of-origin for large patches of adult melanocytes, and two other pigment cell types, iridophores and xanthophores. Hence, Tfap2b confers McSC identity in early development, thereby distinguishing McSCs from other neural crest and pigment cell lineages, and retains multi-fate potential in the adult zebrafish.

**Highlights:**

- Tfap2b and its target genes specify McSCs with mixed pigment cell identities
- Functional dependence on Tfap2b for melanocyte regeneration from the McSC
- *tfap2b* specifies ErbB-dependent McSCs at the stem cell niche
- Fate mapping reveals Tfap2b-McSCs have multi-fate potential for adult pigment cells

## Introduction

Melanocytes produce black-brown pigment that give color and pattern to the animal kingdom, and protect from harmful UV-irradiation (Mort et al., 2015; Nguyen and Fisher 2019). How melanocytes develop in embryogenesis and maintain a regenerative melanocyte population in adulthood is central to our understanding of pigmentation and pattern formation through evolution, as well as in human disease (Baxter and Pavan, 2013; Irion and Nusslein-Volhard, 2019; Kelsh and Barsh, 2011; Kinsler and Larue, 2018; Nusslein-Volhard and Singh, 2017; Parichy and Spiewak, 2015). In melanoma, a deadly cancer of the melanocyte, neural crest and melanocyte developmental mechanisms become reactivated in pathological processes driving melanoma initiation, metastasis, survival and drug resistance (Diener and Sommer, 2020; Johansson et al., 2020; Kaufman et al., 2016; Konieczkowski et al., 2014; Marie et al., 2020; Rambow et al., 2018; Shakhova et al., 2012; Travnickova et al., 2019; Varum et al., 2019; White et al., 2011). Therefore, while mechanisms that underpin neural crest and melanocyte development are important in understanding fundamental cell biology and pattern formation in the animal kingdom, in the disease context, understanding dysregulation and heterogeneity of developmental lineages in cancer provides a rich source of drug targets for the next generation of melanoma therapies (Patton et al., 2021).

Zebrafish are uniquely amenable to advanced imaging of neural crest and pigment cell lineage pathways *in vivo* and have emerged as a powerful model system to study the melanocyte developmental lineage in pattern formation and in melanoma (Kelsh et al., 1996; Owen et al., 2020; Travnickova and Patton, 2021). In zebrafish, embryonic trunk melanocytes are directly derived from the neural crest, but a rare melanocyte subset in the lateral stripe originates from melanocyte stem cells (McSCs; also referred to as MSCs in zebrafish), a postembryonic progenitor population dependent on ErbB-kinase signalling (the epidermal growth factor family of receptor tyrosine kinases). McSCs are essential for replenishing melanocytes following regeneration or during adult growth, and ablation or genetic loss of McSCs leads to large patches of skin devoid of melanocytes cells in the adult fish (Budi et al., 2008, 2011; Dooley et al., 2013; Hultman et al., 2009; Hultman and Johnson, 2010; Taylor et al., 2011). Imaging analysis in zebrafish over time shows that McSCs reside in a niche at the site of the dorsal root ganglia (DRG) and give rise to melanocyte progenitors that migrate along nerves to the epidermis, where they differentiate into pigmented melanocytes (Budi et al., 2011; Dooley et al., 2013; Singh et al., 2016; Singh et al., 2014).

While fate-mapping directly from McSCs has not yet been performed (due to a lack of specific McSC lineage markers), lineage-tracing from the neural crest transcription factor Sox10 proposed that the adult zebrafish pattern is governed by a cell population called MIX cells, a neural crest cell subpopulation that is multi-potent for all three pigment cell types - black-brown <u>m</u>elanocytes, silver iridophores and yellow <u>x</u>anthophores - and that gives rise to neurons and glia (Singh et al., 2016). Adult melanocytes and iridophores are primarily derived from the MIX cells, while adult xanthophores have a dual origin and originate from MIX cells and an embryonic population that expands at the onset of morphogenesis (McMenamin et al., 2014; Singh et al., 2016). While not directly identified, this analysis indicated that MIX cells persist beyond embryonic and larval stages and are present into adulthood, where they are the source of almost all melanocytes that pattern the adult fish (Singh et al., 2016). While it is clear that these pigment MIX cells are derived from the neural crest in zebrafish development, the molecular identification of these cells, how these cells become specified and how they relate to McSCs remains unknown.

Here, through the use of single cell RNA sequencing (scRNA-seq), imaging and lineage tracing approaches, we discover that the transcription factor Tfap2b and a select set of its target genes specify the ErbB-dependent multipotent McSC population. Although mutation in its paralog *tfap2a* is associated with premature hair greying in humans (Praetorius et al., 2013), and results in pigmentation defects in mice and zebrafish (Seberg et al., 2017), the role of *tfap2b* in melanocyte biology is largely unexplored. In melanoma, *TFAP2B* marks a critical node in residual disease cell states (Rambow et al., 2018), although its precise function is yet to be defined. We show that the *tfap2b*-marked cell population resides in the McSC niche and is a progenitor for all three pigment cell lineages. Our data support the conclusion that a Tfap2b transcriptional program specifies a subset of ErbB-dependent cells within a larger embryonic MIX cell population to become McSCs at the dorsal root ganglia. These cells are responsible for generating melanocytes and other pigment cell lineages in adult zebrafish thereby bridging the gap between embryonic and adult patterning.

## Results

### McSCs maintain a neural crest identity at the niche

We performed live imaging of zebrafish embryos to visualize McSCs as they migrate into the DRG-niche site. To this end, we used the transgenic reporter lines *Tg(mitfa:GFP*), that marks melanoblasts, and *Tg(nbt:dsRed*), that marks the neural tube and peripheral nerves (Dooley et al., 2013). Prior to melanization, during the first 22-27 hours post fertilization (hpf), we observed a highly migratory GFP+ subpopulation that traversed the McSC niche and migrated along peripheral nerves to the skin (**Figure 1A, B**). We consider these to be melanocytes that develop directly from the neural crest (NC) and use nerves to migrate to the anatomical location where they generate the embryonic stripes. Subsequently, we observed a distinct, smaller subpopulation of round melanoblasts that was maintained at the niche and a population of stationary columnar cells that coated the nerves (**Figure 1A, C**). These cell populations have previously been described as McSCs and progenitors (Budi et al., 2011; Dooley et al., 2013; Johansson et al., 2020; Kelsh and Barsh, 2011; Singh et al., 2016; Singh et al., 2014). We confirmed that the small, round *mitfa+* cells were McSCs through treatment with an ErbB inhibitor, which eliminated this population at the niche site, as well as stationary nerve-associated progenitors (**Figure 1D-E**) (Budi et al., 2008, 2011; Dooley et al., 2013; Hultman et al., 2009).

**Figure 1:**
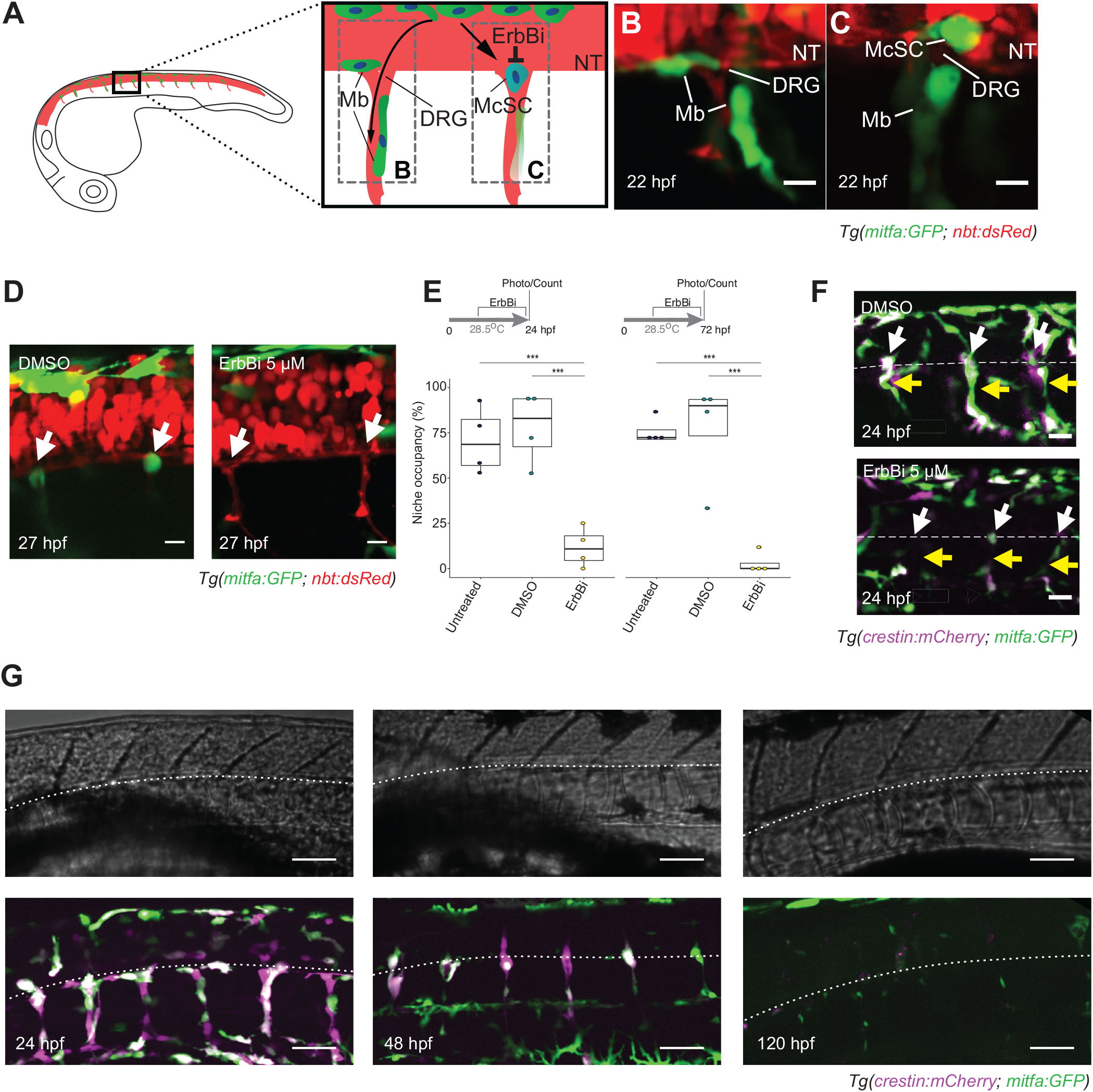
McSCs maintain a neural crest identity at the niche. **A.** Schematic of developing McSC and melanocyte lineages in the zebrafish embryo. *mitfa* expressing melanoblasts (Mbs; green) develop directly from the neural crest and travel to the skin either dorsolateraly (not shown) or ventrally along the neural tube (NT) and peripheral nerves (red; **B**). A subset of those cells, establish at the site of the of the perspective dorsal root ganglia (DRG) and become McSCs (blue cell, **C**). McSCs establishment is sensitive to ErbB-kinase inhibitors (ErbBi). **B.** Melanoblasts migrating along the axons at the neural tube. Confocal stack (20 μm) of *Tg(mitfa:GFP; nbt:dsRed)* embryo imaged laterally at 22 hpf. Scale bar: 20 μm **C.** Newly established McSC at the site of a perspective DRG. Confocal stack (20 μm) of *Tg(mitfa:GFP; nbt:dsRed)* embryo imaged laterally at 22 hpf. Scale bar: 20 μm. **D.** ErbB kinase activity is required for McSCs establishment at the stem cell niche (white arrow). Confocal stacks (20 μm) of *Tg(mitfa:GFP; nbt:dsRed)* embryos treated with either 0.05% DMSO or 5 μM ErbBi. Standard deviation projection. Scale bar: 20 μm. **E.** Quantification of McSC niche occupancy. Tukey-HSD (Honestly Significant Difference) test. ***: p-value <0.0001. **F.** McSCs maintain neural crest identity at the niche. Confocal stacks (30 μm) of *Tg(crestin:mCherry; mitfa:GFP)* embryos treated with either 0.05% DMSO or 5 μM ErbBi. McSCs (white arrows) and nerve-associated cells (yellow arrows) are dependent on ErbB- kinase. Standard deviation projection. Scale bar: 20 μm. **G.** McSCs and nerve associated precursors express *mitfa:GFP* and *crestin:mCherry*, but expression is lost by 120 hpf. Confocal stacks (30 μm) of *Tg(crestin:mCherry; mitfa:GFP)* embryos. The lower edge of the neural tube is indicated (white dotted line) on both fluorescent and corresponding brightfield images (top panels). Brightfield images: average intensity projections; fluorescent images: standard deviation projection. Scale bar: 50 μm.

*Tg(mitfa:GFP*) is expressed by McSCs, but it’s neither an exclusive McSC marker nor does *mitfa* mutation or knockdown prevent acquisition of McSC identity or establishment at the DRG- niche (Budi et al., 2011; Dooley et al., 2013; Johnson et al., 2011), indicating that unknown mechanisms confer McSC identity from the neural crest. To resolve such mechanisms and identify specific McSC markers, we generated a *Tg(crestin:mCherry; mitfa:GFP)* double transgenic line that enabled visualization of both the *mitfa*-expressing melanocyte lineage (GFP+) as well as its origin in the neural crest lineage (mCherry+) (Kaufman et al., 2016) during McSC establishment. In the double transgenic line, we observed newly established McSCs that retained *crestin* expression (GFP+ mCherry+) and cell columns along the peripheral nerves, that were also dependent on ErbB signaling (**Figure 1F**). Notably, however, the expression levels for both GFP and mCherry fluorescence were heterogenous for both cell populations (**Figure 1F-G**). By 48 hpf GFP+ mCherry+ fluorescence was restricted specifically to the McSCs, while differentiating melanoblasts maintained GFP+ fluorescence (**Figure 1G**). In the McSCs, expression of both transgenes was maintained through organogenesis before eventually subsiding by 120 hpf (**Figure 1G**). Thus, unlike differentiating melanoblasts found in the skin, the McSC subpopulation maintains a neural crest identity during the stem cell establishment and specification phase.

### scRNA-seq identifies six distinct neural crest-pigment cell lineage populations

To identify a molecular signature specific for McSCs we needed to first understand how pigment cell populations arise from the neural crest. To this end, we performed droplet-based scRNA-seq on GFP+ and/or mCherry+ cells sorted from the *Tg(crestin:mCherry; mitfa:GFP)* transgenic line using the 10x Chromium system (10x Genomics; **Figure 2A, Figure S1A;** see STAR methods); sequencing 1022 cellular transcriptomes, 996 of which passed our quality control (**Figure S1B-J**). When we visualized the data in two-dimensional space, by applying the Louvain clustering algorithm (Butler et al., 2018) and the Uniform Manifold Approximation and Projection (UMAP; (McInnes et al., 2018) (**Figure 2B**), we found the number of expressed genes was uniformly distributed across the cells (**Figure S1E; Table S1**). To assign cluster identities, we employed a combination of known markers and projections to previously published datasets (Farnsworth et al., 2020; Kiselev et al., 2018; Saunders et al., 2019; Wagner et al., 2018) (**Figure S2C-F, Table S2**).

**Figure 2.**
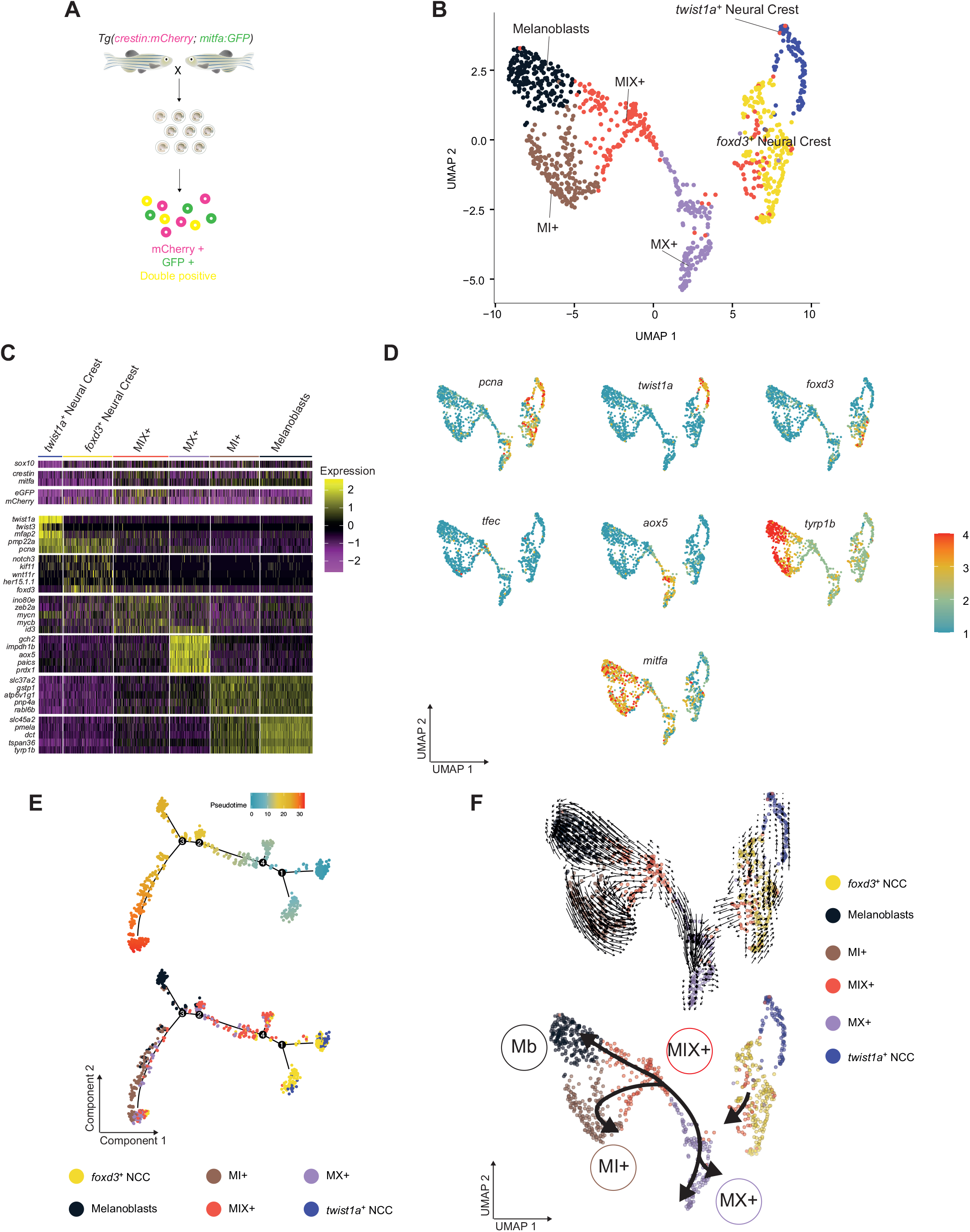
scRNA-seq identifies six distinct neural crest-pigment cell lineage populations. **A.** Schematic of the experimental protocol. Cells that express GFP, mCherry or both were isolated from *Tg(crestin:mCherry; mitfa:GFP)* embryos at 24 hpf. **B.** UMAP of GFP+, mCherry+ and double GFP+ mCherry+ cells (n = 996 cells) obtained from 24 hpf zebrafish embryos after Louvain clustering (dims= 20, resolution = 0.5). **C.** Heatmap showing the average log_2_-fold change expression of five selected genes per cluster identified in **B**. The average log_2_-fold change expression across the 6 clusters of *sox10*, *crestin*, *mitfa*, *mCherry* and *GFP* expression levels are also presented for comparison. **D.** UMAP representations of **B** with color change from blue (negative) to red (positive) based on log_2_ mRNA expression of *pcna*, *twist1a*, *foxd3*, *aox5*, *tfec*, *mitfa*, and *tyrp1b*. **E.** Pseudotime ordering of cells in **B**. Top panel: cells are colored according to their pseudotime scores; bottom panel: cells are colored according to their cluster identity. **F.** RNA velocity analysis of the UMAP represented in **B** (top panel) and simplified representation of the same results (bottom panel).

We identified two proliferative neural crest cell (NCC) populations expressing classical neural crest markers, one of which was characterized by the almost exclusive expression of the bHLH transcription factor and EMT gene *twist1a* (and other *twist* genes; possibly cranial neural crest cells) and the second by expression of *foxd3*, a known Wnt-regulated NC gene (**Figure 2B-D, Table S2**). In addition, we identified four distinct cell clusters that expressed a combination of different chromatophore markers (**Figure 2B-D**). Naming cell clusters from scRNA-seq datasets is challenging because while expression of chromatophore marker genes is indicative of developmental potential, they are not predicative for cell fate; for example *mitfa+* is expressed in cells that differentiate to xanothophores (Parichy et al., 2000). However, we were interested that our scRNA-seq dataset captured cells that are consistent with hypothesized cell populations described through *sox10+* lineage tracing (Singh et al., 2016), gene network studies (Petratou et al., 2021; Petratou et al., 2018), and early imaging studies (Bagnara et al., 1979). One cluster expressed NCC genes, *id* genes, and a chromatophore markers from all three pigment cell types, including melanocyte (*mitfa*, *dct* and *tyrp1b*), iridophore (*pnp4a*) and xanthoblast (*aox5*) markers. For simplicity, we refer to this as a MIX+ cluster because it is positive for expression of markers from all three pigment cell lineages <u>m</u>elanocytes, iridophores and <u>x</u>anthophores. We were intrigued to see that the cells belonging to this cluster expressed less than half of the genes overall. This molecular phenotype is congruent with a stem cell identity, as self-renewing hematopoietic stem cells (HSCs) at the top of the differentiation hierarchy express the lowest number of genes and total mRNA, with total mRNA expression gradually increasing in differentiated cells (Nestorowa et al., 2016) (**Figure S2B**). Cells in the immediately adjacent clusters expressed relatively high levels of melanoblast markers concomitant with markers for two different chromatophores; these included cells that expressed *mitfa* and *aox5* (MX+ cells) and cells that express *mitfa* and *pnp4a* (MI+ cells) (**Figure 2C-D**). Melanoblasts were very similar to MI+ cells in gene expression profile but had a higher expression level of melanocyte differentiation genes. We suggest the melanoblast clusters are the skin-associated *mitfa+* cells that start to be produce melanin by 28 hpf and are responsible for the embryonic melanocyte pattern. We did not find evidence of cells that express iridophore and xanthophore markers without *mitfa+* (IX+ cells) in our experiment.

To understand lineage relationships between the cell clusters, we performed pseudotime analysis (**Figure 2E**). We found the cells to be part of a developmental lineage continuum that originates in the neural crest cell populations and transitions through a MIX+ stage before differentiating into MI+ cells and MX+ cells, or melanoblasts, consistent with a common origin for pigment cells (Bagnara et al., 1979; Petratou et al., 2021; Petratou et al., 2018). A branch point (demarcated as 3) emerged between melanoblasts and the MI+ and MX+ cells, indicative of two distinct melanoblast populations. This is consistent with our imaging analysis (**Figure 1**), which shows *mitfa+* melanoblast populations in the skin (before these become pigmented) and lining the nerves (which are relatively undifferentiated at that stage). Through RNA velocity analysis (La Manno et al., 2018), the developmental lineage relationships were found to be consistent with neural crest cells giving rise to MIX+ cell populations that can either differentiate towards a MX lineage, or initiate expression of melanocyte differentiation genes as either a MI+ or the more differentiated melanoblast cell population (**Figure 2F; Figure S3**). Importantly, despite similar transcriptomes (**Figure 2D**), our analysis indicate that the MI cells are not the progenitors differentiating into melanoblast cells, suggesting that these represent two distinct routes of melanoblast formation in the zebrafish.

### Identification of ErbB-dependent McSCs by scRNA-sequencing

Given that the MIX+ cell population were enriched for *mitfa+ crestin+* double-positive cells (**Figure 2C-D**), we predicted that it could function as a possible source for McSCs. Since McSCs are ErbB kinase dependent, we designed a scRNA-seq experiment in ErbB kinase inhibitor (ErbBi) treated *Tg(crestin:mCherry; mitfa:GFP)* transgenic embryos (**Figure 3A-B**). Clustering 347 cells derived from ErbBi-treated embryos revealed a loss of MI+ cells compared with our untreated dataset. Further, relative to melanoblasts from untreated embryos, the melanoblast population in ErbBi-treated embryos was enriched for cells with reduced differentiation markers, and we therefore called these cells Early melanoblasts (**Figure 3B, Figure S4A**). Pseudotime analysis showed that cells from *Tg(crestin:mCherry; mitfa:GFP)* transgenic embryos form a developmental continuum from NCCs through MIX+ states to MX+ and Early melanoblast populations (**Figure 3C, Figure S4**). Interestingly, in contrast to a loss of cells in the MI+ lineage, *twist1a^+^* NCCs were proportionally increased in ErbBi-treated embryos whereas the early melanoblasts as well as the *twist1a^+^* NCCs were enriched for mesenchymal and proliferative markers including *zeb2a*, *ednrab*, *twist1a*, *pmp22a, pcna* (**Figure 3D-E; Figure S4B-C**).

**Figure 3.**
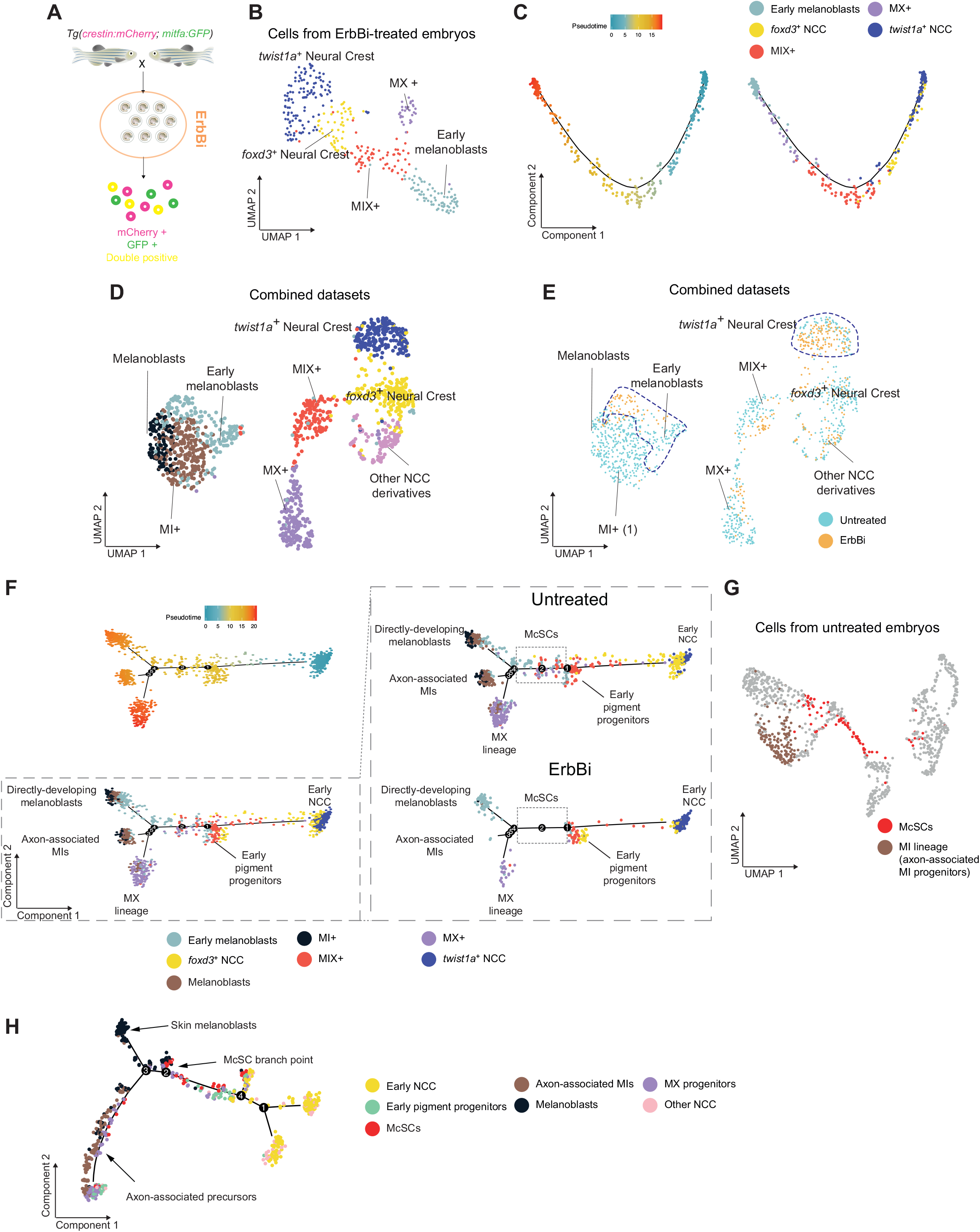
Identification of ErbB-dependent McSCs by scRNA-sequencing. **A.** Schematic of scRNA-seq experimental protocol for ErbBi-treated zebrafish embryos (24 hpf). 5μM ErbBi treatment: 4 - 24hpf. **B.** UMAP of GFP+, mCherry+ and GFP+ mCherry+ double positive cells (n = 346 cells) obtained from ErbBi-treated zebrafish embryos after Louvain clustering (dims = 10, resolution = 0.5). **C.** Pseudotime ordering of cells in **B**. Left panel: cells are colored according to their pseudotime scores; right panel: cells are colored according to their cluster identity. **D.** UMAP of GFP+, mCherry+ and double GFP+ mCherry+ cells (n=1343 cells) from untreated and ErbBi-treated embryos after Louvain clustering (dims = 12, resolution = 1). **E.** UMAP representations of **D** pseudocolored with the origin of the cells analyzed. Dashed lines highlight clusters enriched with ErbBi-treated cells. **F.** Pseudotime ordering of cells in **D**. Cell states present in the untreated embryos (dashed box) are absent in ErbB inhibitor treated embryos. Cells are pseudo-colored according to their pseudotime score (top left panel) or their cluster of origin. Plots in the right panels are split views (by treatment) of the pseudotime analysis. Cell states and the inferred cellular position in the 24 hpf embryo are also indicated. **G.** ErbB-kinase dependent McSCs (red) and the MI progenitors (brown) are highlighted on UMAP presented in **2B**. **H.** Minimum spanning tree presented in **2E** (cells from untreated embryos) pseudocolored according to the cell states described in the lineage trees (**F**). The inferred cellular position and the McSC branch point are indicated.

We conclude from this data, combined with our imaging analysis, that the ErbB-dependent and nerve-associated progenitors represent MI+ cells lost upon ErbBi-treatment (**Figure 3C-E, Figure S4D**). However, from our UMAP, we could not easily distinguish a specific ErbB-dependent MIX+ subpopulation that represent McSCs (**Figure 3E**). To understand the hierarchical dependence of the pigment cell lineage cells emerging from the neural crest, we performed an integrated pseudotime analysis between untreated and ErbBi-treated embryos (**Figure 3F**). In untreated embryos, cells were distributed along a pseudotime continuum from the early NCC to the pigment lineage with early pigment progenitors (node 1) as a “pass through” state. The pigment lineage split into branches ending in defined states for directly developing melanoblasts (node 4), or axon-associated progenitors or the MIX+ population (node 3). When comparing pseudotime ordering of cells derived from ErbBi-treated and untreated embryos, we observed that melanoblast-derived lineages were reduced overall and that axon-associated progenitors, robustly present in untreated embryos were almost entirely lost upon ErbBi treatment. In addition, cells obtained from ErbBi-treated embryos were largely deficient for transitioning cell states (between nodes 1 and 4) but enriched for *twist1a^+^* NCCs (**Figure 3F**), possibly providing an explanation for the reduced number of *foxd3+* NCC populations in the UMAP. Based on these data, we hypothesize that ErbB-dependent cells capable of transitioning between MIX+ and differentiating pigment cell lineages are McSCs. When we mapped these cells back to the untreated *Tg(crestin:mCherry; mitfa:GFP)* analysis, we found that the McSCs formed a subpopulation within the MIX+ cluster (**Figure 3G**), and that the McSCs in fact represent a branch point in the pseudotime analysis (node 2), supporting our hypothesis that their identity is a distinct cell state with the MIX+ cell population (**Figure 3H**).

### McSC identity is specified by a Tfap2b transcriptional program

Next, we performed a differential expression analysis comparing the transcriptomes between the putative McSCs and the combined pigment cell lineages derived from the same wild type embryos (**Figure 4A-B**). As anticipated for a MIX+ subgroup, McSCs expressed all the pigment progenitor markers (**Figure 4A, Table S3**), and combined pigment lineages were enriched for expression of pigment synthesis genes, while the McSCs were enriched for expression of ribosome biogenesis and splicing genes, similar to what has been reported in other stem cell systems (Brombin et al., 2015; Gabut et al., 2020; Gupta and Santoro, 2020; Recher et al., 2013) (**Table S4**). Moreover, McSCs were enriched in genes associated with neurological disabilities in human patients when compared with the early pigment progenitors (**Table S5**).

**Figure 4.**
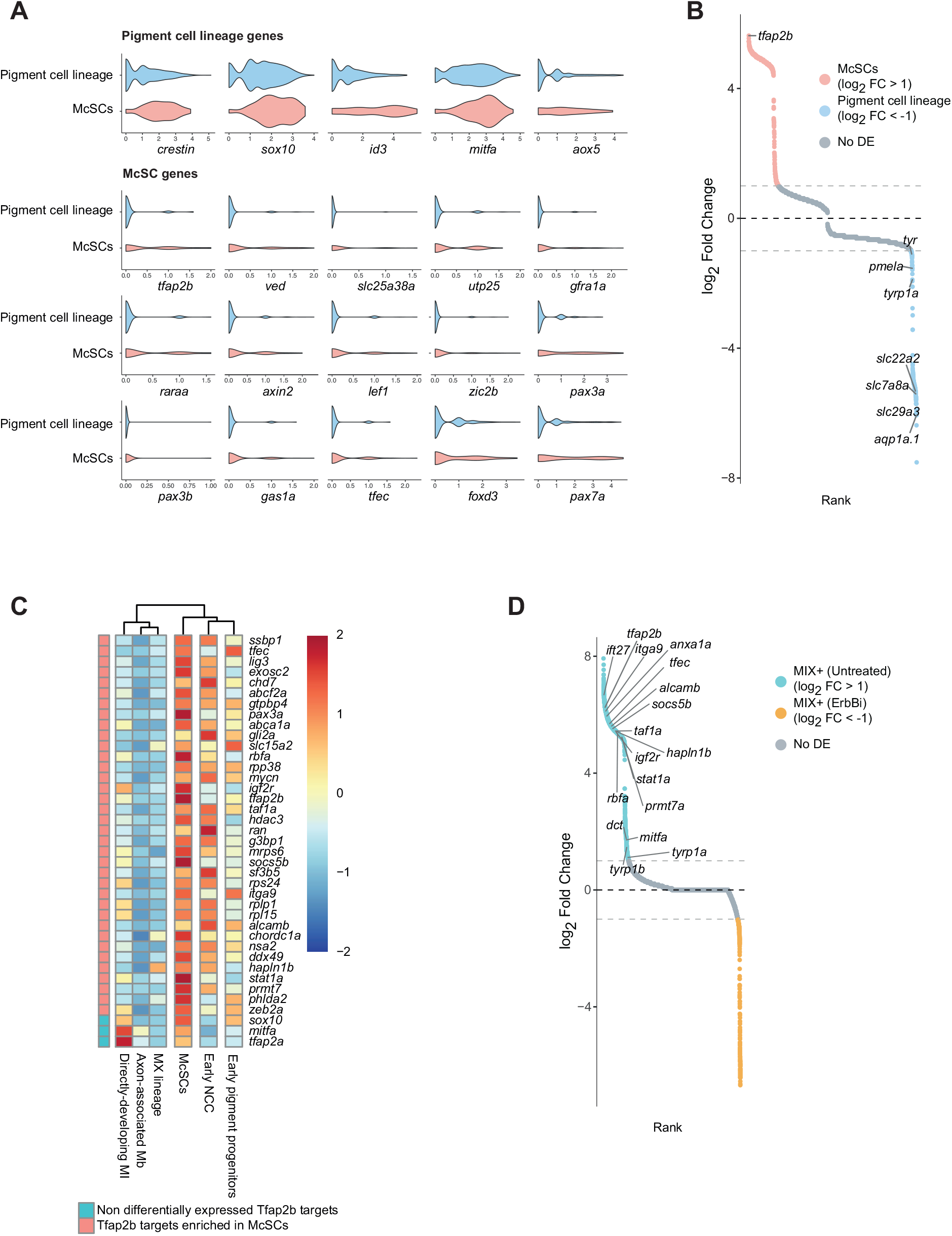
McSC identity is specified by a Tfap2b transcriptional program. **A.** Violin plots of pigment progenitor and McSC gene expression levels. McSCs differentially express a subset of genes (McSC genes) and share expression of genes with cells of the pigment cell lineage (Early pigment progenitors, MI+, MX+ and Melanoblasts from untreated embryos in Figure 3F; **Table S3**). **B.** Rank plot of differential expression analysis between McSCs and all other states of the pigment lineage. The top differentially expressed gene is *tfap2b* (log_2_ fold change: 5.63; adjusted p-value: 4.99e^-5^; **Table S3**). **C.** Clustered heatmap showing the average expression of 36 Tfap2b targets enriched in the McSCs and 3 non-differentially expressed targets (*sox10, mitfa, tfap2a*) (**Table S6**). **D.** Rank plot of differential expression analysis between MIX+ cells between untreated embryos and ErbBi-treated embryos. Most of the McSC-Tfap2b target genes are depleted from MIX+ cells in ErbBi-treated embryos (**Tables S6, S7**).

Strikingly, we found *Transcription Factor AP-2 Beta* (*tfap2b)* to be the top gene specifically enriched within the transcriptome of the putative McSCs (**Figure 4A, B**). Tfap2b is a transcription factor, functionally redundant with Tfap2a, known to regulate neural crest and melanocyte development in mouse, and postulated to differentially specify melanocyte precursors together with Mitfa in zebrafish (Chong-Morrison and Sauka-Spengler, 2021; Lignell et al., 2017; Rothstein and Simoes-Costa, 2020; Seberg et al., 2017). In human melanoma cell xenograft studies, Tfap2b was identified as a marker of a subpopulation present in residual disease states following BRAF plus MEK inhibitor treatment (Rambow et al., 2018), and is also a marker of residual disease in our zebrafish melanoma models (Travnickova et al., 2019) (**Figure S5A**). To investigate whether Tfap2b plays a functional role in regulating the McSCs, we reproduced the Tfap2b (Biotin) ChIP-seq analysis carried out in chicken neural crest tissues (Ling and Sauka-Spengler, 2019). After mapping the chicken Tfap2b targets to their zebrafish homologs, we examined the expression of these genes in McSCs as well as the derivative cell populations within our dataset. We found that a select subset of Tfap2b targets are selectively highly expressed in McSCs compared to other cell states (**Figure 4C**, **Table S6**). Critically, these Tfap2b-McSC target genes are enriched in MIX cells derived from control embryos compared to MIX cells derived from ErbBi-treated embryos (**Figure 4D, Table S7**). These data indicate that Tfap2b plays a pivotal role in the specification of the McSC lineage through activation of a select set of Tfap2b target genes.

### Functional requirement for *tfap2b* in melanocyte regeneration

Next, we asked if *tfap2b* is functionally important for McSC-derived melanocytes but not for melanocytes emerging directly from the neural crest. To answer this question, we used a melanocyte regeneration assay based on a temperature-sensitive *mitfa* mutation, (*mitfa^vc7^*), in which embryos are incapable of generating embryonic melanocytes at higher temperatures, but capable of regenerating melanocytes from McSCs at lower temperatures (Johnson et al., 2011; Zeng et al., 2015). We found that morpholino (MO)-mediated knockdown of *tfap2b* did not impact on the development of most NC-derived melanocytes, but significantly reduced McSC-derived melanocytes in regeneration (**Figure 5A**). These results were confirmed with a second, splicing-site MO knockdown (**Figure 5A**). As McSCs contribute to a small population of melanocytes in the embryonic lateral stripe pattern, McSC activity in zebrafish embryos can be assessed through this as a second independent assay. Indeed, in *tfap2b* knockdown embryos, we found McSC-derived late-stage lateral stripe melanocytes to be reduced, confirming that *tfap2b* is required for McSC-derived melanocyte populations even in non-regenerating embryos (**Figure 5B**). We wanted to assess the impact of *tfap2b* MO knockdown on the adult pigmentation patter, however, although morphants appeared healthy within the first 5 days of development, we were unable to raise the fish beyond 15 days of development, likely due to a potential role for *tfap2b+* cells in other lineages (*e.g.* neural and and craniofacial cells).

**Figure 5.**
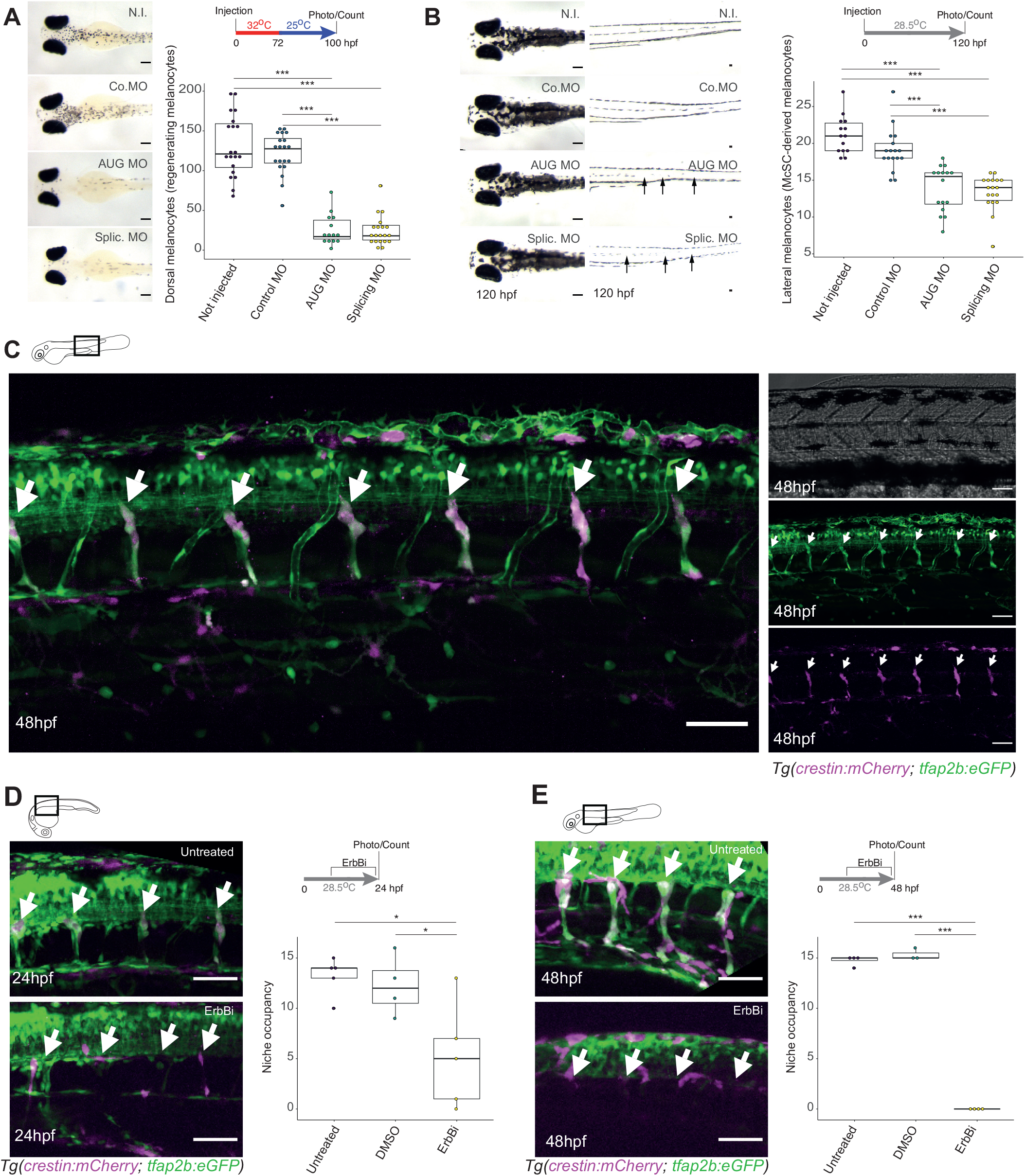
*tfap2b* is expressed at the McSC niche and required for regeneration. **A.** *tfap2b* is required for melanocyte regeneration from the McSC. Images of zebrafish embryos and melanocyte quantification following knock-down of *tfap2b* in a *mitfa^vc7^* regeneration assay. Tukey-HSD (Honestly Significant Difference) test; ***: p-value <0.0001. Scale bar: 200μm. N.I.: Not injected; Co. MO: Control morpholino; AUG MO: AUG-directed morpholino; Splic. MO: Splicing morpholino. Representative of 3 repeated experiments. **B.** *tfap2b* is required for late-stage melanocytes from the McSC. Images of zebrafish embryos and melanocyte quantification following knock-down of *tfap2b*. Only McSC- derived late-developing lateral stripe melanocytes are reduced in *tfap2b* knockdown embryos. Arrows highlight missing lateral stripe melanocytes. Tukey-HSD (Honestly Significant Difference) test; ***: p-value <0.0001. Scale bar: 200μm. Representative of 3 repeated experiments. **C.** Transgenic *tfap2b:GFP* expression in the McSC. Merged image of a double transgenic *Tg(tfap2b:GFP; crestin:mCherry)* zebrafish (left panel) and separated channel images (brightfield, GFP and mCherry channel). White arrows indicate GFP+/mCherry+ double positive McSCs at the DRG. Scale bar: 50μm. **D - E.** *tfap2b+* McSCs require ErbB-kinase at the niche. Tg(*tfap2b:GFP;crestin:mCherry*) embryos at 24 hpf (**D**) and 48 hpf (**E**), either untreated or with 5 μM ErbBi. White arrows indicate McSC niche. Tukey-HSD (Honestly Significant Difference) test; *: p- value=0.0172, ***: p-value <0.0001. Representative of 3 repeated experiments. Confocal stacks (30 μm). Standard deviation projection. Scale bar: 50 μm.

### McSCs are specified by *tfap2b* at the niche

Having established a function for *tfap2b* in McSC-derived regeneration, we next sought to visualize *tfap2b* McSCs during development. To this end, we isolated a 1kb *tfap2b* promoter region and cloned it upstream of the *GFP* coding sequence to generate a transgenic reporter line *Tg(tfap2b:GFP),* and then crossed this line with *Tg(crestin:mCherry)* animals. In double transgenic animals, we identified cells co-expressing GFP and mCherry at the McSC niche and along nerves **(Figure 5C)**. Critically, these cells were absent during ErbBi treatment, indicating that *Tg(tfap2b:GFP)* expression marks the McSC lineage (**Figure 5D-E**).

In extended imaging analysis, we observed that *Tg*(*tfap2b:GFP)* expression peaked during the first 48 hpf but then declined with *crestin:mCherry* expression. We saw no overt evidence of *tfap2b:GFP* expression in adult zebrafish pigment cells, although expression was sustained in the spinal cord in accordance with a neuronal expression pattern (Knight et al., 2005). We did not find evidence of *tfap2b:GFP* expression in other embryonic melanocytes or pigment cells by live imaging, and analysis of our and other datasets confirmed that *tfap2b* expression is principally observed in the McSC population in the pigment cell lineage (**Figure S5B-C**). Together, these results indicate that Tfap2b transiently functions to establish McSC identity during embryogenesis. Thus, Tfap2b plays a role in McSC identity acquisition that is critically required during the first few days of embryonic development.

### *tfap2b-*McSCs have multi-fate potential for all adult pigment cell lineages

MIX+ cells can give rise to the entire complement of pigment cells within a hemisegment (Singh et al., 2016). The highly specific *tfap2b* gene expression in McSCs (**Figure S5B-C**) prompted us to follow the fate of *tfap2b+* cells through development and into the adult pigment pattern. To this end, we cloned the 1kb promoter upstream of *cre* (*tfap2b:cre*) and injected it into the *ubi:switch* transgenic line so that fish were mosaic for *tfap2b:cre* integration and thereby facilitating lineage tracing of cells recombined from GFP+ to mCherry+ (“switched”) cells (Mosimann et al., 2011) (**Figure 6A**). *Tfap2b* is expressed in neural cells in development (Knight et al., 2005), and consistent with this, we found mCherry+ neurons in the dorsal neural tube at 48 hpf (**Figure 6B**, **Figure S5C**). Critically, by 6 days of development, we found mCherry+ McSCs at the niche and in a few melanocytes in the skin. By 13 dpf, we could clearly detect mCherry+ McSCs and nerve-associated cells, as well as a few late-stage embryonic melanocytes in the lateral stripe and xanthophores **(Figure 6C, D)**.

**Figure 6.**
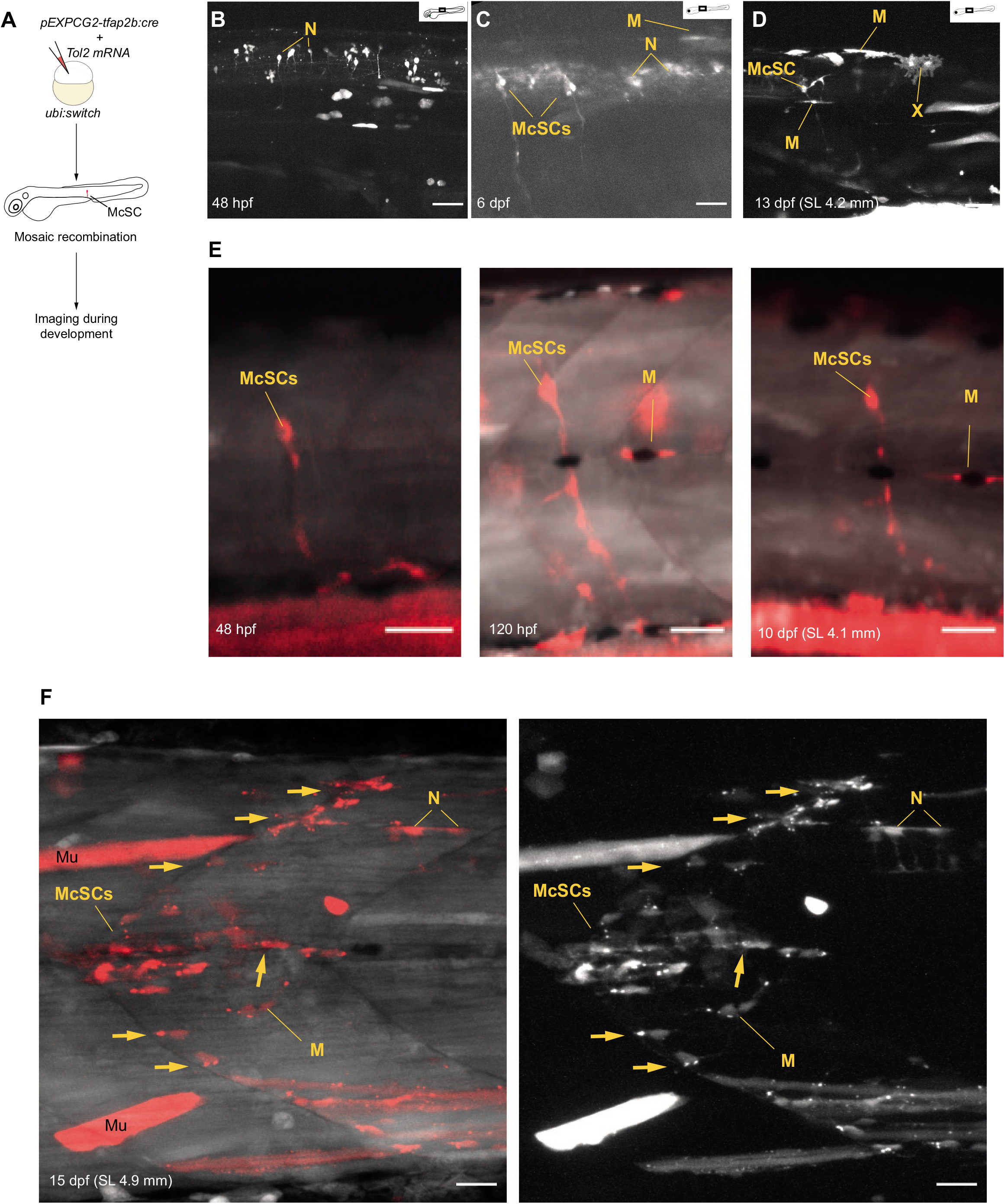
Individual *tfap2b-*McSCs during development and at the onset of metamorphosis. **A.** Experimental protocol overview for *tfap2b+* mosaic lineage tracing. *pEXP-GC2Tol2- tfap2b:cre* and *Tol2* mRNA were injected in *ubi:switch* embryos at the zygote stage and imaged at different stages to identify mCherry expressing McSCs and progenitors. **B - D.** *tfap2b* lineage tracing analysis during zebrafish development. **B:** 48 hpf, **C:** 6 dpf, **D**: 13 dpf (SL 4.2 mm). Representative images of 7-8 animals imaged per stage. White pseudo-coloring is used for the mCherry channel. M: melanocytes; McSCs: melanocyte stem cell; N: neural tube neurons; SL: standard length; X: Xanthophore. Maximum Intensity projection. Scale bars: 50μm. **E.** Example of a single McSC followed through the development of a single fish and imaged at 48 hpf (left panel), 120 hpf (middle panel) and 10 dpf (SL 4.1 mm). The McSC produces progenitors that populate the ventrally located axonal projection and a lateral line- associated melanocyte (M). Embryo injected with 1.5 pg/nl of *tfap2b:cre*. Scale bars: 50μm. **F.** Dramatic expansion of progenitors from the McSCs at the onset of the metamorphosis. Confocal imaging of a clone expanding from a single McSC at 15 dpf (SL 4.9 mm). Arrows indicate axon-associated cells. M: melanocyte, Mu: muscle (off-target), N: spinal cord neuron. White pseudo-coloring is used for the GFP channel and red is used for the mCherry channel in the left panel, white pseudo-coloring is used for the mCherry channel in the right panel. Embryo injected with 1.5 pg/nl of *tfap2b:cre*. Scale bars: 20μm.

To follow the fate of *tfap2b+* McSCs over time, we first needed to be confident that the tracing of cell fates and clones originated from a single or a few recombination events. To this end, we injected limited dilutions of *tfap2b:cre* (1.5 and 3.125 pg/nl), and analyzed 70 embryos at 2 dpf: 54 embryos expressed no mCherry signal at the McSC, but expressed mCherry in other cells (*e.g.* the neural tube); 13 expressed mCherry at a single visible McSC; and three embryos expressed mCherry at a few McSCs (with lower level of certainty due to overlap with neuronal cells by epifluorescence microscopy). This indicates that these experiments are within the limiting transgene range and that mCherry+ cells or clones at later stages of development are likely the result of a single or a few recombination events in the McSCs. We followed these fish over 5 days, and could clearly visualize cells emerging from the McSC along nerves. In **Figure 6E**, we show an example of an McSC giving rise to cells along nerves and a late-stage lateral line melanocyte by 5 and 10 dpf. In another example shown in **Figure 6F**, we detected an expansion of mCherry+ cells emerging from the McSC site at the onset of metamorphosis (15 dpf), including a newly differentiating melanocyte.

To understand the fate of *tfap2b+* cells in the adult pigmentation pattern, we analyzed 56 zebrafish at one month of age by confocal imaging. By one month, mCherry+ melanocytes, xanthophores and iridophores were particularly prominent in the developing stripes during metamorphosis (**Figure 7A-B**), and occasionally nerves. We identified MIX clones that spanned the whole dorso-ventral axis in hemisegments and were clearly visible in the scales. We observed one fish with a sparse clone made up of only a few iridophores, xanothophores and melanocytes distributed along the length of the hemisegment and not adjacent to each other (**Figure 7C**). This may indicate that McSCs do not all have equal potential to generate progenitors or may represent a late-stage McSC activation event.

**Figure 7.**
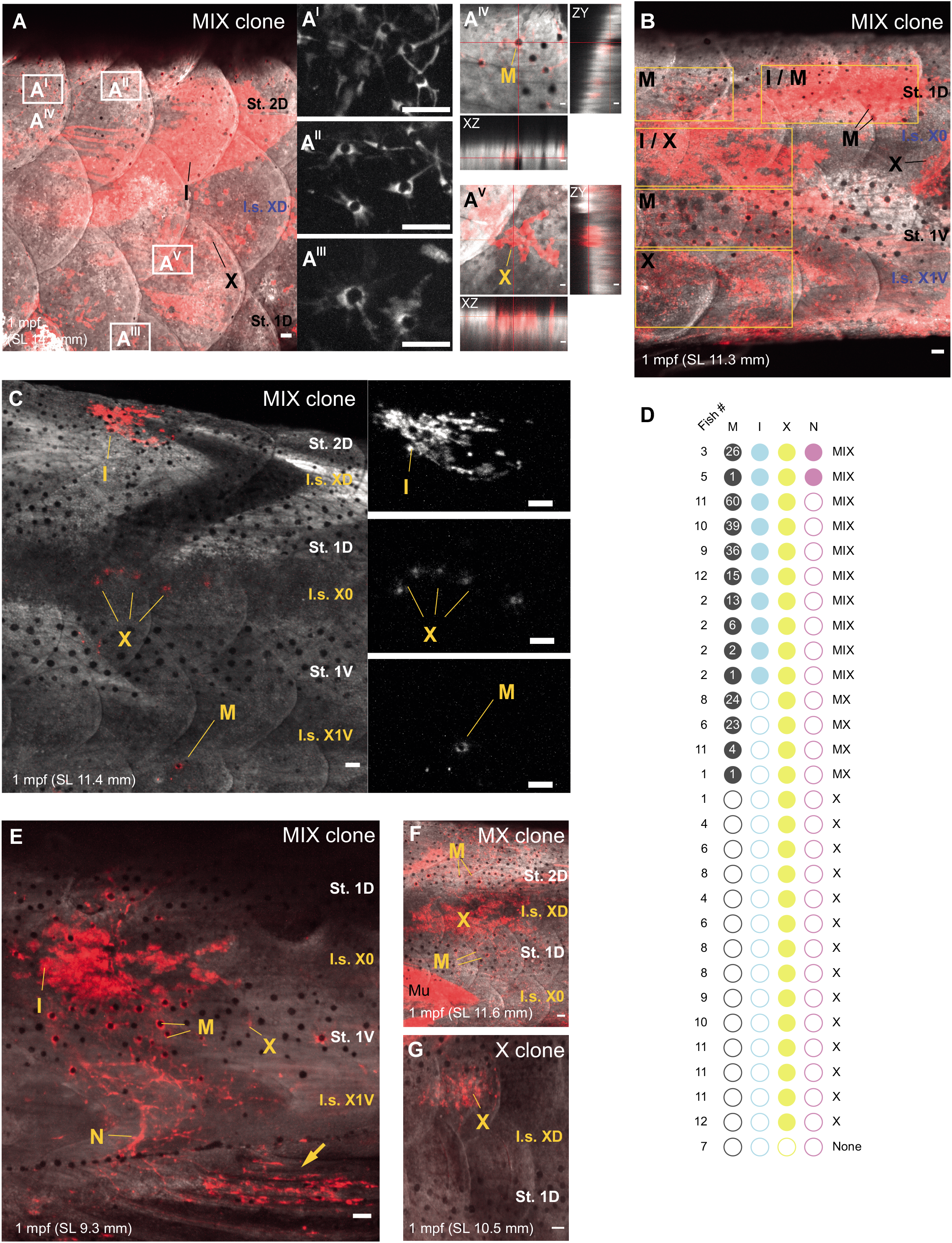
*tfap2b-*McSCs have multi-fate potential for all adult pigment cell lineages. **A - C.** MIX clones in *tfap2b* lineage tracing analysis in the adult pigment pattern. **A:** caudal trunk; **B:** tail region; **C**: medial trunk. Magnified images of melanocytes are presented in stripe 2D (**A^I^**, **A^II^**), and in stripe 1D (**A^III^**). Orthogonal projections of a melanocyte (**A^IV^**) and a xanthophore (**A^V^**) showing that cells are localized in the scale. **B** shows a MIX clone spanning the dorso-ventral axis. **C** depicts a sparse MIX clone, with magnifications of iridophores, xanthophores and a melanocyte (from top to bottom). Pseudo-coloring is used for the GFP channel and red is used for the mCherry channel in **A-C**; white pseudo-coloring is used for the mCherry channel in magnified panels and **A^I-III^** and in **C** magnified images. Representative images of >20 fish, each injected with 25 pg/nl of *tfap2b:cre*. I: Iridophore, M: melanocyte, X: xanthophore. I.s.: Interstripe, St.:Stripe. Maximum Intensity projection. Scale bars: 50 μm. **D.** Frequencies of the different derivatives in clearly defined clones in 12 juvenile *ubi:switch* zebrafish injected low doses of *tfap2b:cre* plasmid (1.5 pg/nl, 3.25 pg/nl, 6 pg/nl). I: Iridophore, M: melanocyte, N: clone-associated nerve, X: xanthophore. Clones were named according to the pigment cell composition. Numbers in the black dot represent mCherry+ melanocytes within the clone. **E - G.** Representative images of clones analyzed in **D**. **E:** MIX clone with a clone-associated nerve on the caudal trunk; **F:** MX clone on the medial trunk; **G**: X clone on the rostral trunk. **E** shows a MIX clone spanning the dorso-ventral axis with a clone-associated nerve. The arrow indicates clone extension into the anal fin. I: Iridophore, M: melanocyte, Mu: muscle (off target signal), X: xanthophore. I.s.: Interstripe, St.:Stripe. Maximum Intensity projection. Scale bars: 50 μm. All fish are derived from embryos injected with 1.5 pg/nl of *tfap2b:cre*.

Our experimental cohort of zebrafish included fish injected with a range of *tfap2b:cre* concentrations (1.5 pg/nl to 25 pg/nl). We observed the same patterns of labelled cells in the fish injected at all concentrations of the transgene, however, some very large clones spanned one or more segments along the rostro-caudal axis indicating that the Cre recombination might have occurred in multiple consecutive McSCs. Therefore, we quantitated the clones derived from 12 fish at one month of age that had been injected with low concentrations of *tfap2b:cre* (1.5 pg/nl, 3.25 pg/nl, 6 pg/nl), and all had clearly definable clone boundaries. Clone numbers on these fish ranged from 0 - 5 clones / fish, but on average had 2 clones per animal. Even at this low *tfap2b:cre* concentration, many fish had large, clearly visible MIX clones that filled the hemisegment (**Figure 7D**). Some MIX clones also contained nerves (as seen by Singh *et al*.) and even expanded into the fin regions, as seen in **Figure 7E**. Melanocyte numbers per clone varied from 1 to 60. We also observed large clones composed of just melanocytes and xanthophores (MX clones, **Figure 7F**), as well as a number of smaller clones within the interstripes derived of a single pigment cell type, the xanthophores (X clones, **Figure 7G**). Once again, these data suggest that McSCs may not have equal potential to generate all derivatives. Single xanthophore clones could also arise from a *tfap2b+* McSC derived xanthophore (as seen in **Figure 6D** at 6 dpf), or alternatively, from a separate, as yet unknown, *tfap2b+* xanthoblast progenitor at later developmental stages. Finally, we performed cryo-sectioning to analyze *tfap2b+* derived cells within the 1 month zebrafish, and found mCherry+ cells migrating along nerve tracks, as well in the skin and in the scales **(Supplementary Figure S6)**. Together, these findings confirm that *tfap2b* labels McSC populations at the niche, and that these cells have a multi-potent identity with the potential to give rise to all three adult pigment cell types.

## Discussion

McSCs provide an on-demand source of melanocytes during growth, metamorphosis and regeneration, however, we have lacked an understanding of how McSC populations are specified from the neural crest or melanocyte progenitors in embryogenesis. Here, we discover that Tfap2b and a select set of its target genes specify the McSC from the neural crest, and that these cells reside at the DRG niche. Molecularly, our data points to a population of cells with mixed pigment cell identity (MIX+ cells), only some of which activate a Tfap2b transcription program to become the ErbB-dependent McSCs at the niche. We show Tfap2b is required for melanocyte regeneration from the McSC, providing a functional role for Tfap2b in stem cell potential. Moreover, we show that Tfap2b-McSCs, that are established early in development, are fated to contribute to all three pigment cell lineages in the adult zebrafish. Conceptually, our data support a new model in which Tfap2b expression in the neural crest leads to activation of a select set of target genes within a broader MIX cell population to specify the McSCs. Once specified, these cells have the potential to generate all three pigment cell types in the adult.

scRNA-sequencing combined with live imaging reveals that the spatiotemporal development of pigment cell fates is complex. Molecularly, our study resolves two developmental pathways for melanocytes; melanoblasts, that develop directly from the neural crest, and MI+ cells, that are ErbB-signalling dependent. We propose that the melanoblasts contribute to the embryonic pattern in the skin, while the MI+ cells are the ErbB-dependent *crestin+ mitfa+* cells that line the peripheral nerves and remain relatively undifferentiated, as seen in **Figure 1**. We also find the McSC transcriptomes cluster within a larger MIX+ cell population, and propose that not all MIX+ cells in embryogenesis have the potential to contribute to the adult pattern. While the McSCs are ErbB-dependent, the ErbB-independent MIX+ cells may represent a tri-potent precursor cell for the three chromatopohore lineages in the embryo pigment pattern (Bagnara et al., 1979; Petratou et al., 2021; Petratou et al., 2018). Tfap2b and its transcriptional program would thereby distinguish McSCs from other MIX+ cells.

Despite the strong functional evidence for McSC activity at the DRG, the lack of cell type-specific markers have prevented investigations into molecular mechanisms underpinning their biology. Indeed, additional McSC populations may be present in the zebrafish embryo that are dependent on other microenvironment niches, including an ErbB-dependent and blood vessel-associated population dependent on Endothelin factors (Camargo-Sosa et al., 2019). Our lineage tracing analysis proves that *tfap2b-*McSCs are multi-potent and give rise to melanocytes, iridophores and xanthophores of the adult. These findings provide a foundation for studying how McSCs make fate decisions in growth and replenish pigment cells in during tissue regeneration.

Understanding how McSCs are specified and activated in normal development, and how they become dysregulated in disease, is critical for regenerative medicine and cancer biology. In mammals, multiple McSC populations have been identified at distinct anatomical locations. In the skin, a melanocyte stem cell population residing in the hair follicle is a reservoir for hair and skin melanocytes during the hair cycle and for re-pigmentation in vitiligo, and is a cellular origin of melanoma (Lee and Fisher, 2014; Moon et al., 2017; Sun et al., 2019). In the hair follicle during the resting phase (telogen), McSC populations are found at the bulge region and the secondary hair germ, and are functionally heterogeneous (Joshi et al., 2018; Joshi et al., 2019). While both cell populations have melanocyte potential, bulge McSCs are CD34+ and have myelination potential, while the secondary hair germ McSCs are CD34- and have enhanced melanocyte regeneration potential. Notably, RNA-seq analysis from isolated McSCs subpopulations show that CD34+ McSCs express higher levels of neural crest stem cell genes (*e.g. Ngfr, Twist1/2, Snai2, Sox9*) while CD34- MsSCs express higher levels of melanocyte transcriptional network genes (*e.g. Sox10, Mitf, Erbb3, Tyr, Tyrp1*), including *Tfap2a* and *Tfap2b*. On hairless skin, such as the palm or sole, the sweat gland serves as a niche for melanocyte - melanoma precursors (Okamoto et al., 2014). In the dermis, a multi-potent stem cell that expresses neural-crest markers NGFRp75 and Nestin is a source of extrafollicular epidermal melanocytes, as well as mesenchymal and neuronal lineages (Li et al., 2010; Zabierowski et al., 2011). These cell populations may be similar to the multi-potent NC-derived Schwann-cell precursor (SCP) population in chick and mouse that reside along the growing nerve, representing a niche for various cell types, including melanocytes (Adameyko et al., 2009; Diener and Sommer, 2020; Ernfors, 2010; Furlan and Adameyko, 2018). Zebrafish do not have hair, but the zebrafish McSC anatomical niche suggests that zebrafish McSCs are functionally analogous to the SCP population found in birds and mammals (Budi et al., 2011; Dooley et al., 2013). Our findings that Tfap2b and a select set of target genes specifically marks the McSCs at the DRG open new doors to address how nerves provide a niche for melanocyte progenitors (Furlan and Adameyko, 2018). Further, given the enriched *Tfap2b* expression in CD34- McSCs, our results may have relevance for the how hair follicle McSCs selectively contribute to melanocyte regeneration (Joshi et al., 2019).

As one of the most aggressive and heterogenous cancers, the melanoma transcriptional landscape spans developmental neural crest and melanocyte lineage signatures, stem cell signatures, and trans-differentiation signatures (Diener and Sommer, 2020; Marine et al., 2020; Patton et al., 2021). We propose that the molecular mechanisms that regulate McSC biology have direct relevance to melanoma pathogenesis. Illustrating this, we recently discovered that the rate of differentiation from the McSC lineage in zebrafish is dependent on a PRL3-DDX21 transcriptional elongation checkpoint and that this same mechanism in melanocyte regeneration portends poor outcomes for melanoma patients (Johansson et al., 2020). Importantly, we and others find that Tfap2b is expressed in human and zebrafish melanoma residual disease cell states, a malignant and drug-resistant cell state that contributes to disease recurrence (Marine et al., 2020; Rambow et al., 2018; Shen et al., 2020; Travnickova et al., 2019) (**Figure S5A**). Thus, the developmental Tfap2b mechanism we identify here for zebrafish McSCs could be co-opted in melanoma, such that Tfap2b melanoma residual disease cell states may represent a dysregulated McSC developmental lineage.

## Author Contributions

Conceptualization: EEP, AB; Methodology: AB, JT; Software: AB, DJS, JT, YL; Validation: AB, HB, ZZ; Formal analysis: AB, DJS, YL; Investigation: AB, JT, HB, ZZ, AIJY; Resources: EEP; Writing original draft: EEP, AB; Writing review and editing: All authors; Visualisation: AB, AIJY; Supervision: TC, EEP; Funding acquisition: TC, EEP.

## Acknowledgements

We are grateful to Cameron Wyatt and the IGC Zebrafish Facility for zebrafish management and husbandry, Elisabeth Freyer and the IGC FACS facility, Ann Wheeler and the IGC Imaging Facility for supporting the imaging experiments, Edinburgh Genomics for sequencing, Christina Lilliehook for editing support, and Lauren Saunders and David Parichy for helpful discussions on scRNA-sequencing analysis. DJS is funded by the Medical Research Council (Doctoral Training Programme in Precision Medicine). TC is funded through a Chancellor’s fellowship held at the University of Edinburgh. EEP is funded by MRC HGU Programme (MC_UU_00007/9), the European Research Council (ZF-MEL-CHEMBIO-648489), and Melanoma Research Alliance (687306).

## KEY RESOURCES TABLE

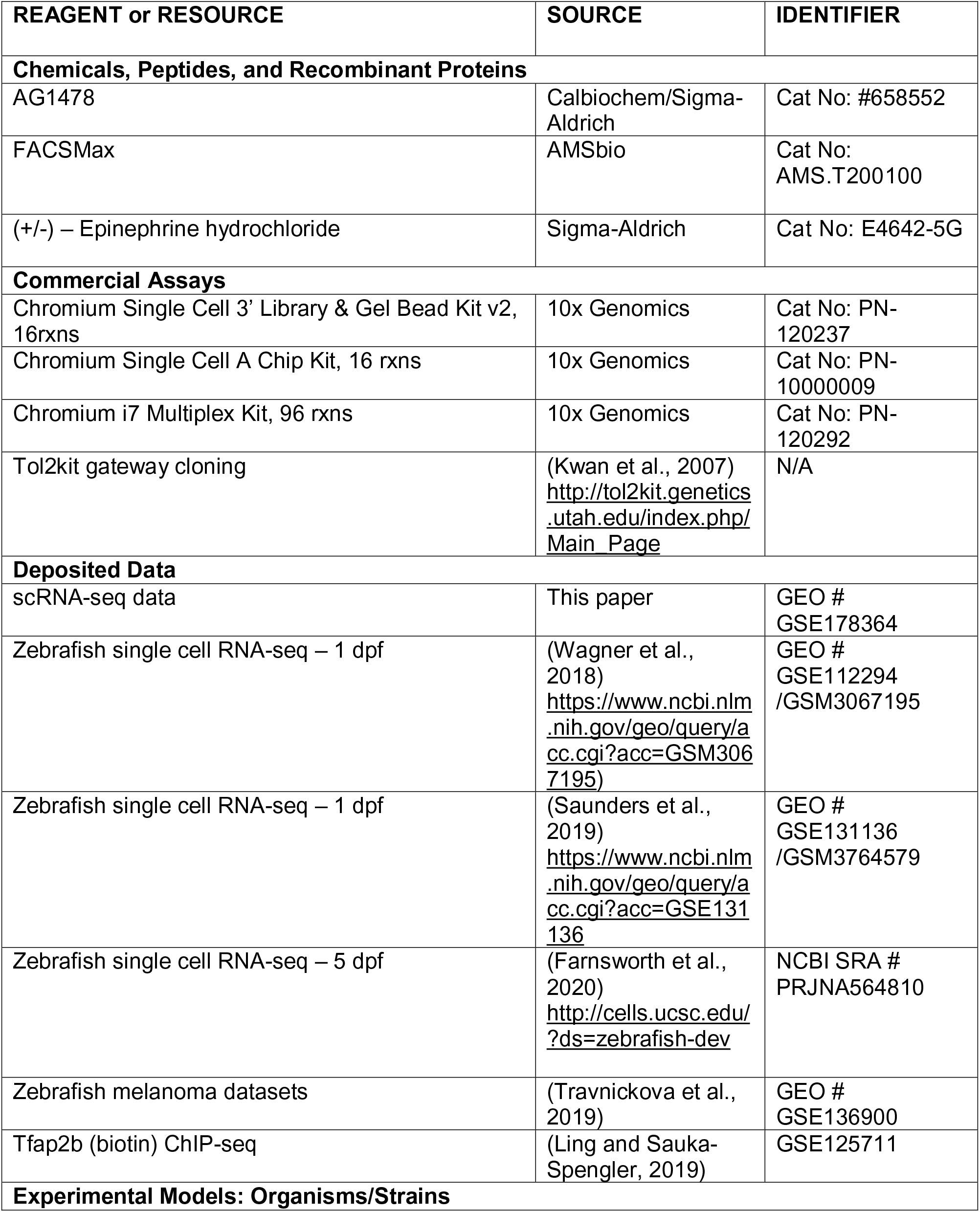
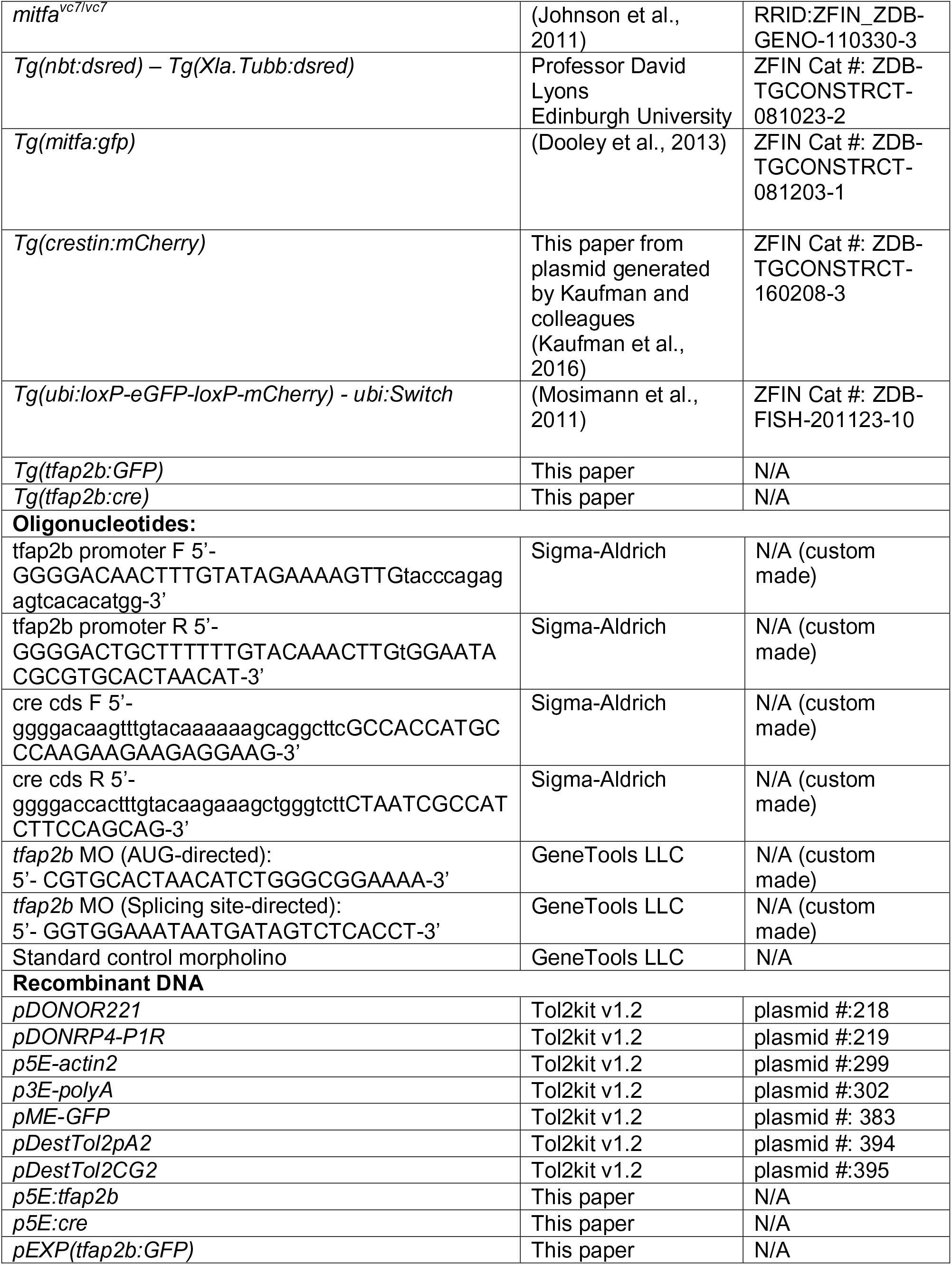
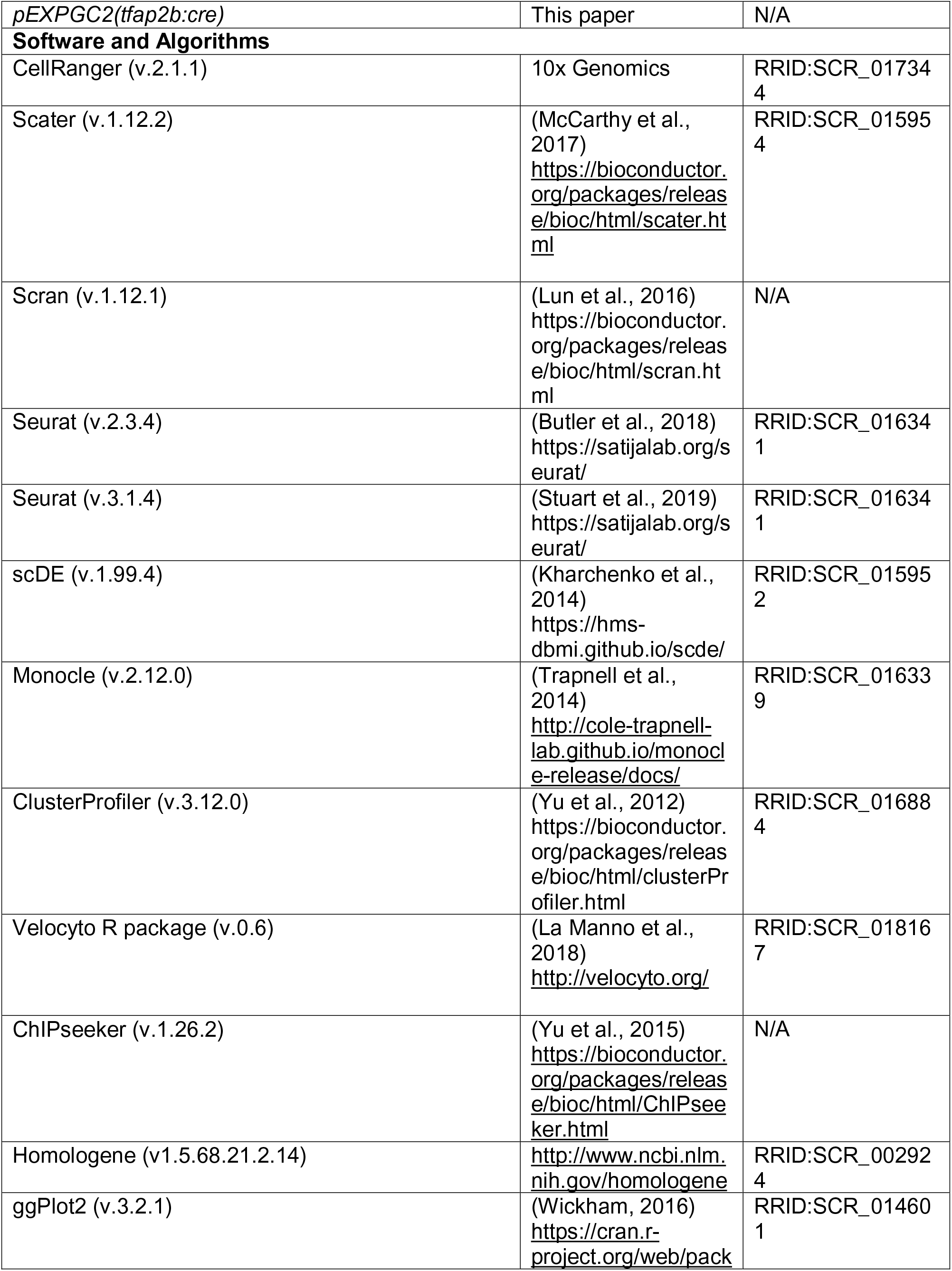
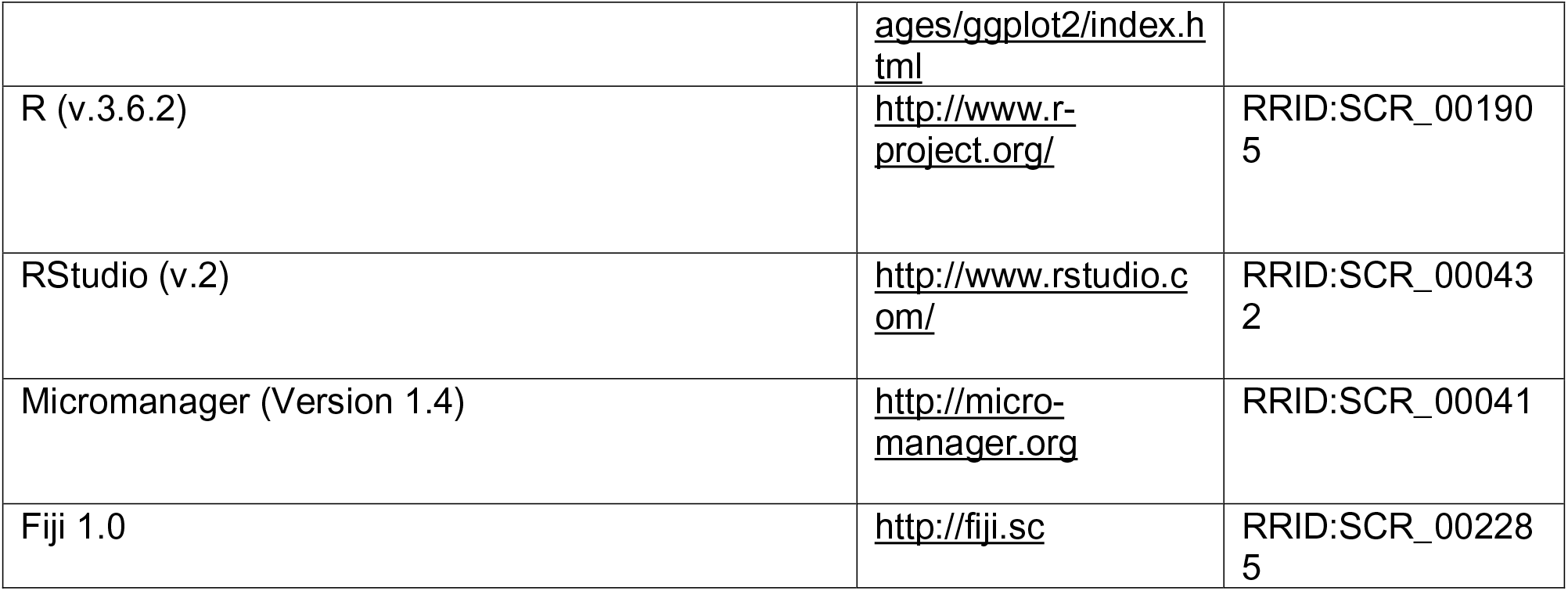

## STAR METHODS

### RESOURCE AVAILABILITY

#### Lead contact

Further information and requests for resources and reagents should be directed to and will be fulfilled by the Lead Contact, E. Elizabeth Patton.

#### Materials availability

Newly generated materials from this study are available upon request to the Lead Contact, E. Elizabeth Patton.

#### Data and code availability

scRNA-seq experiments have been submitted to GEO: GEO # GSE178364. All other data and codes supporting the findings of this study are available from the Lead contact upon reasonable request.

### EXPERIMENTAL MODELS

Zebrafish were maintained in accordance with UK Home Office regulations, UK Animals (Scientific Procedures) Act 1986, amended in 2013, and European Directive 2010/63/EU under project license 70/8000 and P8F7F7E52. All experiments were approved by the Home office and AWERB (University of Edinburgh Ethics Committee).

Fish stocks used were: wild-type AB, *mitfa^vc7^* (Johnson et al., 2011; Zeng et al., 2015), *Tg(nbt:dsRed), Tg(mitfa:GFP)* (Dooley et al., 2013), *Tg(crestin:mCherry)* generated for this study from the plasmid kindly provided from Charles Kaufman (Washington University), *Tg(ubi:loxP-GFP-loxP-mCherry; ubi:Switch*) (Mosimann et al., 2011), *Tg(tfap2b:GFP)* (this study). Combined transgenic and mutant lines were generated by crossing. Adult fish were maintained at ∼28.5°C under 14:10 light:dark cycles. Embryos were kept at either 25°C, 28.5°C or 32°C and staged according to the reference table provided by Kimmel and colleagues (Kimmel et al., 1995) or Parichy and colleagues (Parichy et al., 2009)

## METHODS

### Generation of zebrafish transgenic lines

The promoter region of *tfap2b* (1kb upstream of the first coding exon) was PCR amplified from zebrafish total genomic DNA with the following set of primers: forward: 5’- GGGGACAACTTTGTATAGAAAAGTTGtacccagagagtcacacatgg-3’; reverse: 5’- GGGGACTGCTTTTTTGTACAAACTTGtGGAATACGCGTGCACTAACAT-3’. Amplicons were cloned into *pDONRP4-P1R* (Tol2Kit v1.2, plasmid #: 219) to generate the *p5E-tfap2b* entry clone which were combined with either *pME-GFP* (Tol2Kit v1.2, plasmid #: 383) (Kwan et al., 2007), and the SV40 polyA sequence from *p3E-polyA* (Tol2Kit v1.2, plasmid #: 302) into the *pDestTol2pA2* (Tol2Kit v1.2, plasmid #: 394) destination vector to generate *pEXP(tfap2b:GFP)* and expression vectors.

The cre coding sequence was amplified from *pMC-CreN* plasmid (kind gift Jianguo Shi Check) with the following primers: forward: 5’- ggggacaagtttgtacaaaaaagcaggcttcGCCACCATGCCCAAGAAGAAGAGGAAG-3’; reverse 5’- ggggaccactttgtacaagaaagctgggtcttCTAATCGCCATCTTCCAGCAG-3’. The amplicon was cloned into *pDONOR221*(Thermo Fisher) producing a middle entry vector, *pME-cre* that was cloned with the *tfap2b* promoter from *pME-tfap2b*, and the SV40 polyA sequence from p3E- polyA into the *pDestTol2CG2* destination vector (Tol2Kit v1.2, plasmid #: 395) to generate the *pEXPGC2(tfap2b:cre)* expression vector.

The pEXP vectors were mixed with *Tol2* mRNA (in vitro transcribed with the Ambion mMessage mMachine SP6 Kit, Thermo Fisher, from the Tol2Kit *pCS2FA-transposase* plasmid – plasmid #: 396); and microinjected into 1-cell stage either AB or *ubi:switch* embryos, at a final concentration 25 pg/nl and 35 pg/nl respectively. In order to perform lineage trancing with a sufficiently limiting amount of transgene, we also injected *pEXPGC2(tfap2b:cre)* expression vector at decreasing concentrations (12.5 pg/nl, 6.25 pg/nl, 3.125 pg/nl, or 1.5 pg/nl) with 35 pg/nl of Tol2 transposase.

Zebrafish embryos expressing the *tfap2b:GFP* transgene were selected and grown to adulthood before crossing with wildtype zebrafish to obtain the F1 generation. Embryos expressing the *tfap2b:cre* transgene were selected and either grown until the desired developmental stage for confocal imaging or single housed for imaging in epifluorescence.

### Zebrafish morpholino oligonucleotides

For the *tfap2b* morpholino, 1ng of AUG-directed morpholino (5’- CGTGCACTAACATCTGGGCGGAAAA-3’) or 2ng of splicing morpholino (5’- GGTGGAAATAATGATAGTCTCACCT-3’) oligonucleotide were injected, as well as the standard control (Gene Tools, LLC). Regenerating melanocytes in *tfap2b* morpholino-injected and control-injected fish were tested in the *mitfa^vc7^* regeneration assays with embryos raised at 32°C for 72h before down-shifting to 25°C. Imaging and quantification of regenerating melanocytes were performed at 120 hpf. Late-developing melanocytes in *tfap2b* morpholino-injected and control-injected fish were tested during normal development in AB embryos raised at 28.5°C for 120h before imaging and quantification of regenerating melanocytes. Representative of 3 biological replicates.

### Cryosections

*mCherry+ ubi:switch* juveniles were cut in the middle of a clearly visible clone before being fixed in 4% PFA overnight at 4°C. The fixed juvenile fish were then soaked in 15% sucrose/PBS overnight at 4°C, then 30% sucrose/PBS for 6 hours at 4°C prior to being embedded in O.C.T compound (Sakura) and frozen in isopentane. 40μm cryosections were cut using a Leica cryostat.

### Imaging

Embryos at 4 hpf *Tg(mitfa:GFP; nbt:dsRED)*, *Tg(crestin:mCherry; mitfa:GFP) or Tg(tfap2b:GFP; crestin:mCherry)* were arrayed in 6-well plates (Corning) containing 0.05% DMSO or 5 μM AG1478 (ErbB-inhibitor, ErbBi, Sigma-Aldrich) in 3 ml of E3 embryo medium and kept at 28°C until imaging time. *ubi:swit*ch embryos injected with *pEXPGC2(tfap2b:cre)* were screened under a fluorescence stereomicroscope for the presence of GFP in the heart at 48 hpf and were then raised as described previously. The same microscope was used to follow 70 embryos injected with either 3.125 or 1.5 pg/nl of *pEXPGC2(tfap2b:cre)* plasmid. Repeated imaging was performed at 48 hpf, 120 hpf, 10 dpf and 15 dpf. Larvae older than 4 dpf were first soaked in 5 mg/ml epinephrine (Sigma-Adrich) for 5 minutes, anestethized and quickly mounted in 3% methyl-cellulose prior imaging.

Embryos or fish were selected randomly for confocal imaging as above. Fish older than 5 dpf were imaged while in terminal anaesthesia. 1 mpf fish were first soaked in 5mg/ml epinephrine (Sigma-Adrich) for 10 minutes prior mounting in low-melting point agarose. Images of randomly picked embryos or selected adult fish were acquired using a 0.4x/0.3, a 10X/0.5 or a 20X/0.75 lens on the multimodal Imaging Platform Dragonfly (Andor technologies, Belfast UK) equipped with 405, 445, 488, 514, 561, 640 and 680nm lasers built on a Nikon Eclipse Ti-E inverted microscope body with Perfect focus system (Nikon Instruments, Japan). Data were collected in Spinning Disk 40μm pinhole mode on the Zyla 4.2 sCMOS camera using a Bin of 1 and no frame averaging using Andor Fusion acquisition software. If required: Z stacks were collected using the Nikon TiE focus drive.

Images of cryosections were acquired using 10 and 20X Lenses on a Zeiss Axio-Observer Z1 inverted microscope (Carl Zeiss UK, Cambridge, UK), with a ASI MS-2000 XY stage (Applied Scientific Instrumentation, Eugene, OR). Samples were illuminated using Brightfield or a Lumencor Spectra X LED light source (Lumencor Inc, Beaverton, OR) complete with Chroma #89000ET single excitation and emission filters (Chroma Technology Corp., Rockingham, VT) and acquired on either a Prime BSI Express camera for fluorescence microscopy or a QImaging Retiga R6 Colour (Teledyne Photometrics) for colour brightfield imaging.

For melanocyte counting, regenerating and normal developing embryos were fixed in 4% PFA/PBS and dehydrated in increasing concentrations of glycerol (Sigma-Adrich). Images were acquired Leica MZFLIII fluorescence stereo microscope with a 1x objective fitted with a

Qimaging Retiga Exi CCD camera (Qimaging, Surrey, BC, Canada). Image capture was performed using Micromanager (Version 1.4).

Data were analyzed using Fiji 1.0 and 64bit Java8. Representative of 3 biological repeats.

### Single cell suspensions, fluorescence activated cell sorting and library preparation

*Tg(crestin:mCherry; mitfa:GFP)* were processed in two instances and the following method applied to each treatment separately to obtain two libraries.

300 – 400 embryos at 4 hpf were divided in two equally sized batches and arrayed in 6-well plates (Corning) containing either E3 or 5 μM AG1478 (ErbBi) in 3 ml of E3 till 24 hpf. A single cell suspension of each batch of embryos was then produced following the method described by Manoli and colleagues (Manoli and Driever, 2012) with minor modifications. Samples were sorted by a FACSAria2 SORP instrument (BD Biosciences UK). Green fluorescence was detected using GFP filters 525/50 BP and 488 nm laser, red fluorescence was detected using RFP filters 582/15 BP and 561 nm laser, and live cells selected with DAPI using DAPI filters 450/20 BP and 405 nm laser. Prior to sorting for fluorescence levels, single cells were isolated by sequentially gating cells according to their SSC-A vs. FSC-A and FSC-H vs FSC-W profiles according to standard flow cytometry practices. Cells with high levels of DAPI staining were excluded as dead or damaged. Cells from wild-type stage matched embryos (without transgenes) were used as negative control to determine gates for detection of mCherry and GFP fluorescence. Then *Tg(crestin:mCherry; mitfa:GFP)* cells from either untreated or ErbBi-treated zebrafish were purified according to these gates. 10,000 (AB background) fluorescent cells per batch were collected in 100μl of 0.04% Bovine Serum Albumin (BSA)/PBS in LoBind tubes (Fisher Scientific), spun down at 300G at 4°C, resuspended in 34μl of 0.04% BSA/PBS, and immediately processed using the Chromium platform (10x Genomics) with one lane per sample. Single-cell mRNA libraries were prepared using the single-cell 3’ solution V2 kit (10x Genomics). Quality control and quantification assays were performed using High Sensitivity DNA kits on a Bioanalyzer (Agilent).

Libraries were sequenced on an Illumina NovaSeq platform (1 lane of a S2 flowcell, read 1: 26 cycles, i7 Index: 8 cycles, read 2: 91 cycles). Each sample was sequenced to an average depth of at least 1750 million total reads. This resulted in an average read depth of ∼50,000 reads/cell after read-depth normalisation.

## QUANTIFICATION AND STATISTICAL ANALYSIS

Statistical details of the experiments, n numbers, and dispersion and precision measurements can be found in the figure legends.

### Bioinformatics analysis

#### scRNA-seq data processing and quality check

FastQ files were aligned using the CellRanger (v.2.1.1, 10x Genomics) pipeline to custom zebrafish STAR genome index using gene annotations from Ensembl GRCz11 release 94 with manually annotated entries for *GFP*, *mCherry*, *mitfa* intron 5 and *mitfa* intron 6 transcripts, filtered for protein-coding genes (with Cell Ranger *mkgtf* and *mkref* options). Final cellular barcodes and UMIs were determined using Cell Ranger. Libraries were aggregated (using 10X Cell Ranger pipeline ‘cellranger aggr’ option), with intermediary depth normalization to generate a gene-barcode matrix.

Gene-cell matrices (total: 1519, from untreated embryos: 1022, from ErbBi-treated embryos: 497) were uploaded on R (v.3.6.2) and standard quality control metrics with the Scater package (v.1.12.2)(McCarthy et al., 2017). Only cells with total features >700, log10 total counts > 3.0, and mitochondrial gene counts (%) < 10 were considered as high quality and kept for further analyses (total:1343, from untreated embryos: 996, from ErbBi-treated embryos: 347). Prediction of the cell cycle phase was performed using the cyclone function in the Scran (v.1.12.1) (Lun et al., 2016).

### Clustering, UMAP visualisation and cluster calling

The Louvain clustering of the separated libraries (**Figure 2B, 3B**) was performed with Seurat (v.3.1.4) (Stuart et al., 2019) using the *FindNeighbors* and *FindClusters* functions (cells from untreated embryos: dims = 20, resolution = 0.5; cells from ErbBi-treated embryos: dims = 10, resolution = 0.5) after performing linear dimensionality reduction and checking the dimensionalities of the datasets visualized with elbow plots.

Data were projected onto 2 dimensional spaces using Uniform Manifold Approximation and Projection (UMAP) (McInnes et al., 2018) using the same dimensionality values listed above.

Cluster specific genes were identified using the *FindAllMarkers* and *FindMarkers* function in Seurat (v.2.3.4 or v.3.1.4) with default parameters (Wilcoxon Rank Sum test that compares a single cluster against the others) and then using a Bayesian approach with the scDE package (v.1.99.4) (Kharchenko et al., 2014). See **Table S2**.

Cluster calling was done after detection of published marker genes for specific cell types and by making unbiased pairwise comparisons based on gene overdispersion against published datasets GEO #: GSE112294 (Wagner et al., 2018), GEO #: GSE131136 (Saunders et al., 2019), and NCBI SRA #: PRNJNA56410 (Farnsworth et al., 2020) using the scMap package (v.1.6.0) (Kiselev et al., 2018) and between the datasets presented in this paper.

The combined clustering for the aggregated libraries used for McSC identification (**Figure 3D-E**, **S4B-D**) was performed using the Seurat package (v.2.3.4, dims=12, resolution=1) (Butler et al., 2018).

Plots were generated either using Seurat (v.3.1.4) or ggplot2 (v.3.2.1) (Wickham, 2016).

### Pseudotime analyses and comparison of the developmental lineages

Differential expression analyses were performed using the identified clusters within each dataset to resolve pseudotemporal trajectories using the *setOrderFilter*, *reduceDimension* and *orderCells* function in Monocle (v.2.12.0) (Trapnell et al., 2014). The minimum spanning tree obtained from cells derived from the untreated or ErbBi embryos was rooted on cluster “*twist1a^+^* Neural Crest” (**Figure 2F**, **3C** and **3H**).

The integrated pseudotime analysis used for the discovery of the McSCs (**Figure 3F**) was based on the combined clustering (**Figure 3D-E**). The cluster “Other NCC derivatives” was excluded from this analysis because it did not contain pigment cell markers, and we wanted to understand the development of the pigment cell lineage. The top 1000 highly dispersed genes among the untreated embryos dataset were chosen as feature genes to resolve the lineage tree using the *setOrderingFilter*, *reduceDimension*, and *orderCells* functions of Monocle (v.2.12.0). We used the default parameters (except for max_components = 4 and norm_method = log) to generate the 3D trajectory during dimensionality reduction. The same genes were then used to order the cells from the ErbBi-treated embryos and then the combined results were plotted using the *PlotComplexTrajectories* function and highlight missing states (states 7,8,11 that were collectively called “McSCs” and plotted back in the original UMAPs) in the ErbBi-treated dataset. The transcriptome of the cells belonging to the McSC states from untreated embryos were then compared with the ones from cells of the states composing the pigment lineage (“Axon-associated MIs”, “Directly-developing melanoblasts ”, “MX lineage” and “Pigment progenitors”). The differential expression analysis was performed using a Bayesian approach with the scDE package (v.1.99.4) and the results were plotted using the ggplot2 package (v.3.2.1) and the pathway analysis was performed using the ClusterProfiler R package (v.3.12.0). The same approach was used for the other differential expression analyses presented.

### RNA velocity analyses

RNA velocity analyses were performed with the Velocyto R package (v.0.6) (La Manno et al., 2018) using default parameters.

### Tfap2b targets

The Tfap2b (Biotin) ChIP-seq data was retrieved from the Gene Expression Omnibus (GEO) under the accession code GSE125711. The same mapping and peak calling pipeline described by the dataset-linked publication was used, with the setting of FDR < 0.01, fold enrichment > 2 set to define the final Tfap2b binding element region. R Package “ChIPseeker” (v1.26.2) (Yu et al., 2015) was used to map the peak coordinates to the gene symbols using chicken genome assembly galGal5. Package “homologene” (v1.5.68.21.2.14) was used to identify human homologous for the gene targets identified in Tfap2b (Biotin) ChIP-seq, as well as the differential expressed genes in the zebrafish cluster (with human homolog *ZEB2* manually mapped to zebrafish gene *zeb2a* as we found it was not automatically mapped by the software).

### Other statistical analyses

Counts of dorsal melanocytes in the head and trunk region were performed using the Cell Counter plugin on ImageJ Fiji.

The niche occupancy was calculated as the percentage of GFP+ positive cells per number of visible DRG (**Figure 1**) or as the number of fluorescent cells ventrally to the neural tube (**Figure 5**). Two images per embryos were acquired (caudally to the first somite, ∼8 somites; rostrally to the urogenital opening, ∼8 somites) and the total number of the niches per embryo were considered.

Statistics for regeneration assays and niche occupancy were performed using running R (v.3.6.2) from RStudio (v.2). For all assays, a normal distribution and equal variance were assumed. For assays with more than two groups, data was analyzed through Analysis of variance (ANOVA), using Tukey-HSD (Honestly Significant Difference) test. Box plots: boxes represent 25^th^ to 75^th^ percentiles, lines are plotted at median. Whiskers represent Min to Max.

## SUPPLEMENTAL INFORMATION

### Supplementary Figure Legends

**Figure S1.**
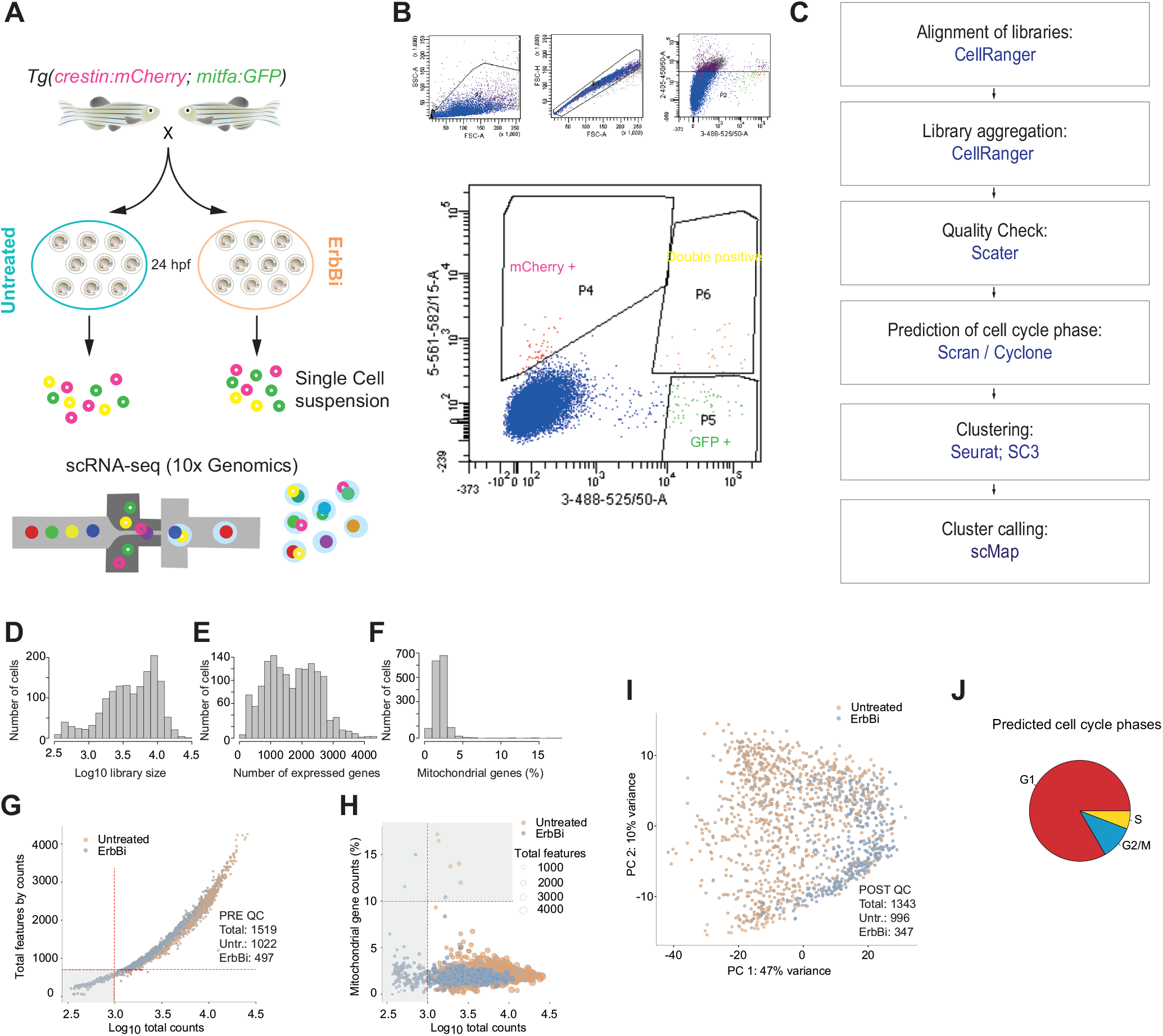
Experimental design, analysis pipeline, and quality check results for scRNA-seq experiments. **A.** General schematic of the experimental protocol. Collected embryos were separated into two equally sized batches, and one half reared in standard E3 and the other half treated with 5μM AG1478 (ErbB inhibitor, ErbBi) between 6 and 24 hpf. Single cell suspensions were generated from each batch of embryos and only DAPI-, GFP+ and/or mCherry+ were collected and processed with the 10x Genomics Chromium system. **B.** Representative FACS plots of the sorted populations. **C.** Summary of the analysis pipeline followed. **D - F.** Bar plots showing the relation to the Log_10_ library size (**D**), the number of expressed genes (**E**), and the percentage of mitochondrial genes (**F**) of the aggregated libraries from cells pre-quality check (QC). **G - H.** Scatter plots distribution of the total features (genes) expressed in relation to the Log_10_ total counts (**G**) and the percentage of mitochondrial genes in relation Log_10_ total counts (**H**) of the aggregated libraries from embryos pre-QC. Red dashed lines indicate the QC thresholds (Total features by counts > 700; Log_10_ total counts > 3.0; Percentage of mitochondrial genes <10%). Gray areas highlight the cells that were considered low quality and excluded from downstream analyses. Orange: cells from untreated embryos, Blue: cells from ErbBi-treated embryos. **I.** Principal component analysis (PCA) of the aggregated libraries from cells post-QC. Orange: cells from untreated embryos, Blue: cells from ErbBi-treated embryos. **J.** Pie chart of the proportion of cells in a predicted cell cycle phase within the aggregated libraries from cells post-QC.

**Figure S2.**
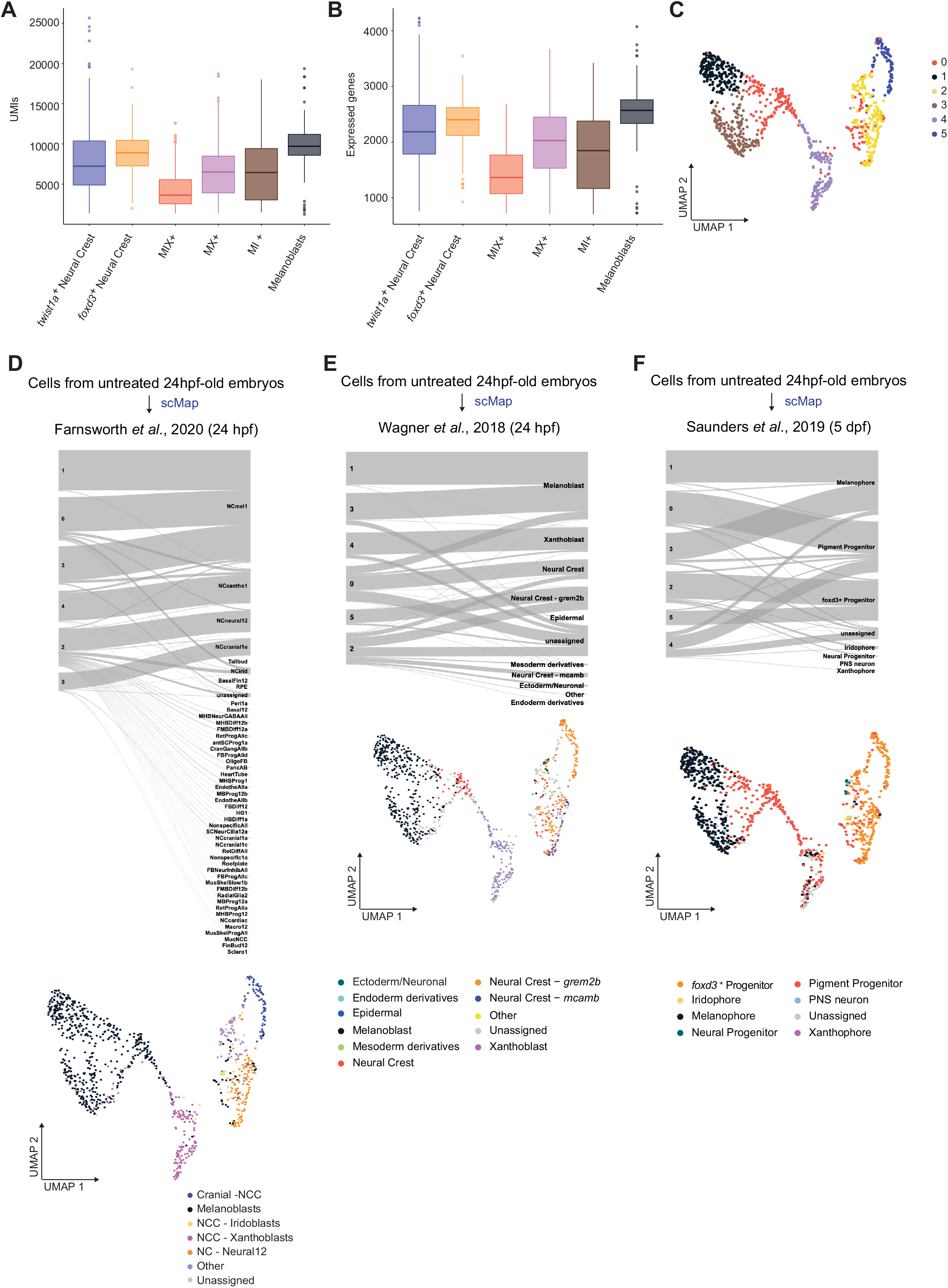
Proportion of expressed genes and comparison with known datasets. **A, B.** Distribution of the Unique Molecular Identifiers (UMIs, **A**) and the expressed genes (**B**) in each cell of the dataset in the different clusters. **C.** UMAP of GFP+/DAPI-, mCherry+/DAPI-, and GFP+/mCherry+/DAPI- cells (n = 996 cells) obtained from untreated 24 hpf zebrafish embryos after Louvain clustering (dims= 20, resolution = 0.5) showing 6 clusters. The original clusters identifiers are indicated. **D.** SanKey plot and UMAP representation based on the scMap results obtained by comparison of the untreated dataset and the Farnsworth *et al*., 24 hpf dataset. **E.** SanKey plot and UMAP representation based on the scMap results obtained by comparison of the untreated dataset and the Wagner *et al*., 24 hpf dataset. **F.** SanKey plot and UMAP representation based on the scMap results obtained by comparison with the Saunders *et al*., *sox10:cre+* 5 dpf zebrafish embryos dataset.

**Figure S3.**
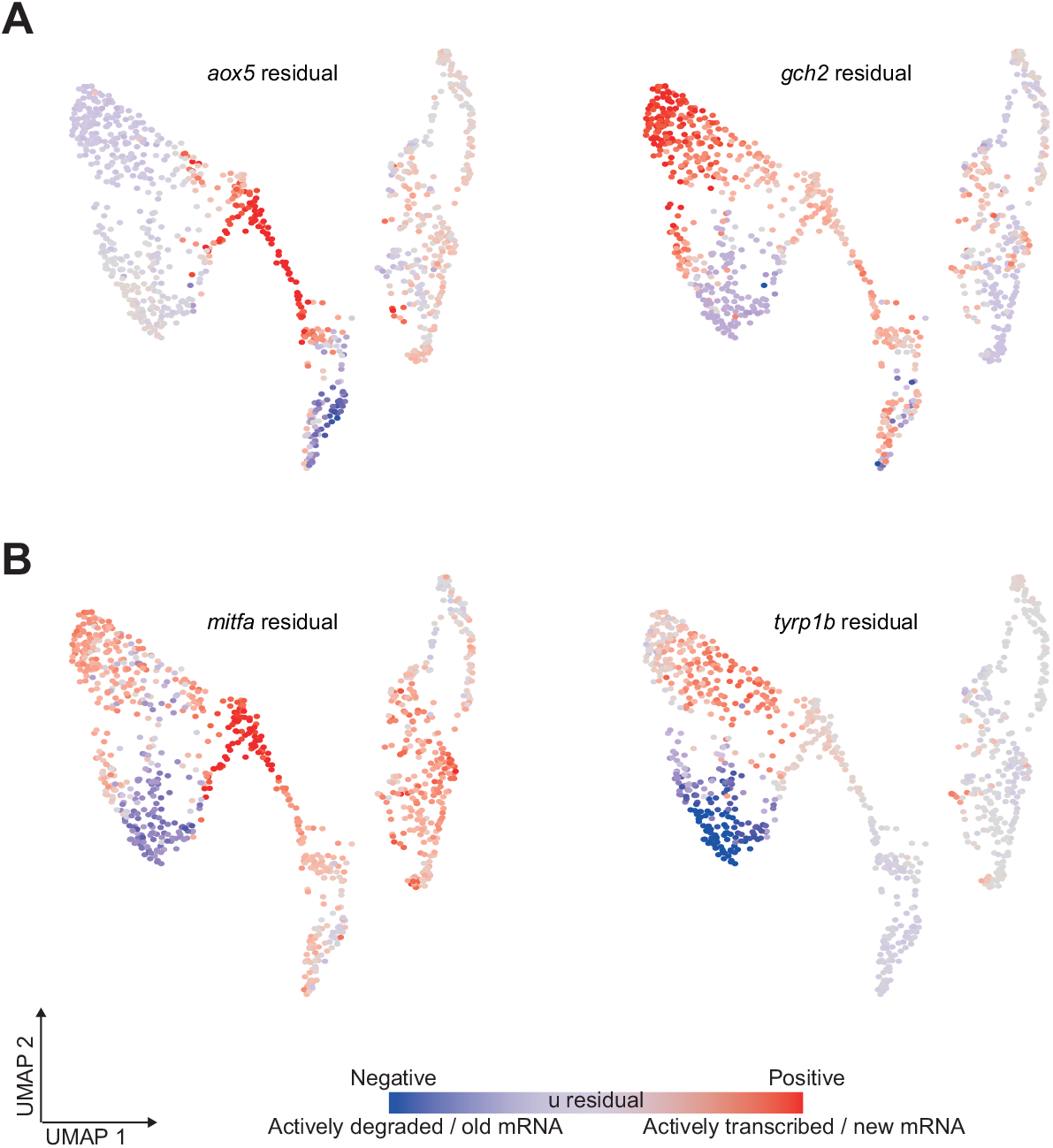
mRNA splicing (RNA velocity) of lineage markers. UMAP representations of Figure 2B with ratios of actively transcribed and degraded transcripts of selected genes. Read counts were pooled across the five nearest cell neighbors.

**Figure S4.**
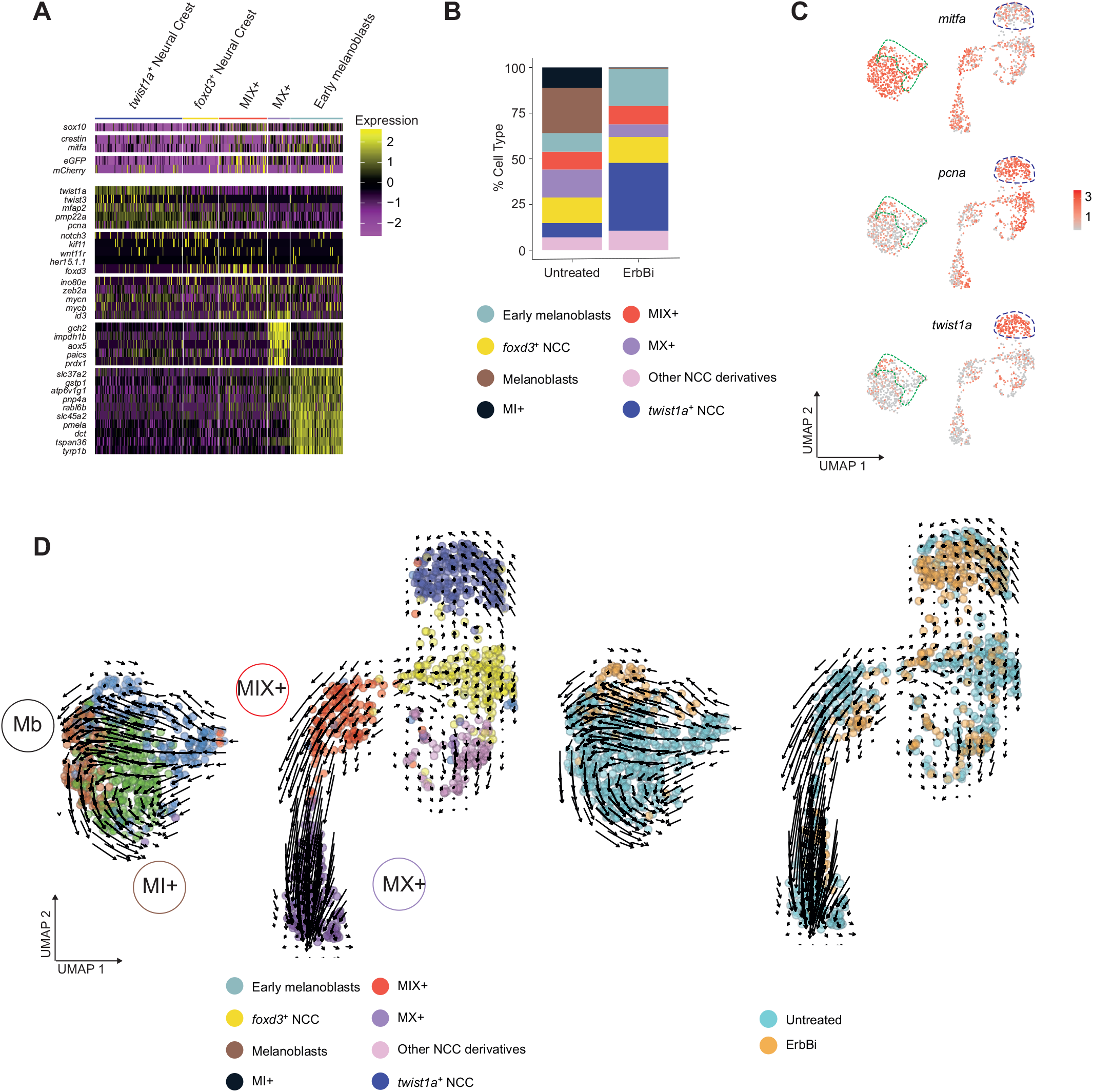
ErbB-kinase supports melanocyte lineage differentiation. **A.** Heatmap showing the average log_2_-fold change expression of five selected genes per cluster identified in Figure 3B. **B.** Bar plot representing the differences in the proportions of cells within each cluster from untreated and ErbBi-treated embryos. Cells derived from ErbBi-treated and untreated embryos contribute differently to most of the clusters: Other NCC derivatives, p-value = 0.04153; “*twist1a^+^* Neural Crest”, p-value < 2.2e-16; “Early melanoblasts”, p-value = 2.195e^- 6^; MX progenitors, p-value = 9.123e^-5^; MI progenitor, p-value <2.2e^-16^; Melanoblasts, p-value = 4.988e^-10^). The proportions of cells in the *foxd3^+^* Neural Crest (p-value=1) and the MIX progenitors (p-value = 0.9342) clusters are not statistically different. 2-sample test for equality of proportions with continuity correction (Z-score test). **C.** UMAP representations of Figures 3D and 3E with color change from grey (negative) to red (positive) based on log_2_ mRNA expression of *mitfa, pcna and twist1a*. Cells derived from ErbBi-treated embryos in the “Early melanoblasts” cluster (green dashed line) express genes which are expressed by melanoblasts and *twist1a^+^* Neural Crest (blue dashed line). **D.** RNA velocity analysis reported on the UMAPs represented in Figure 3D (left panel) and Figure 3E (right panel). M: melanoblasts; MI: melano-iridophore state; MX: melano- xanthophore state.

**Figure S5.**
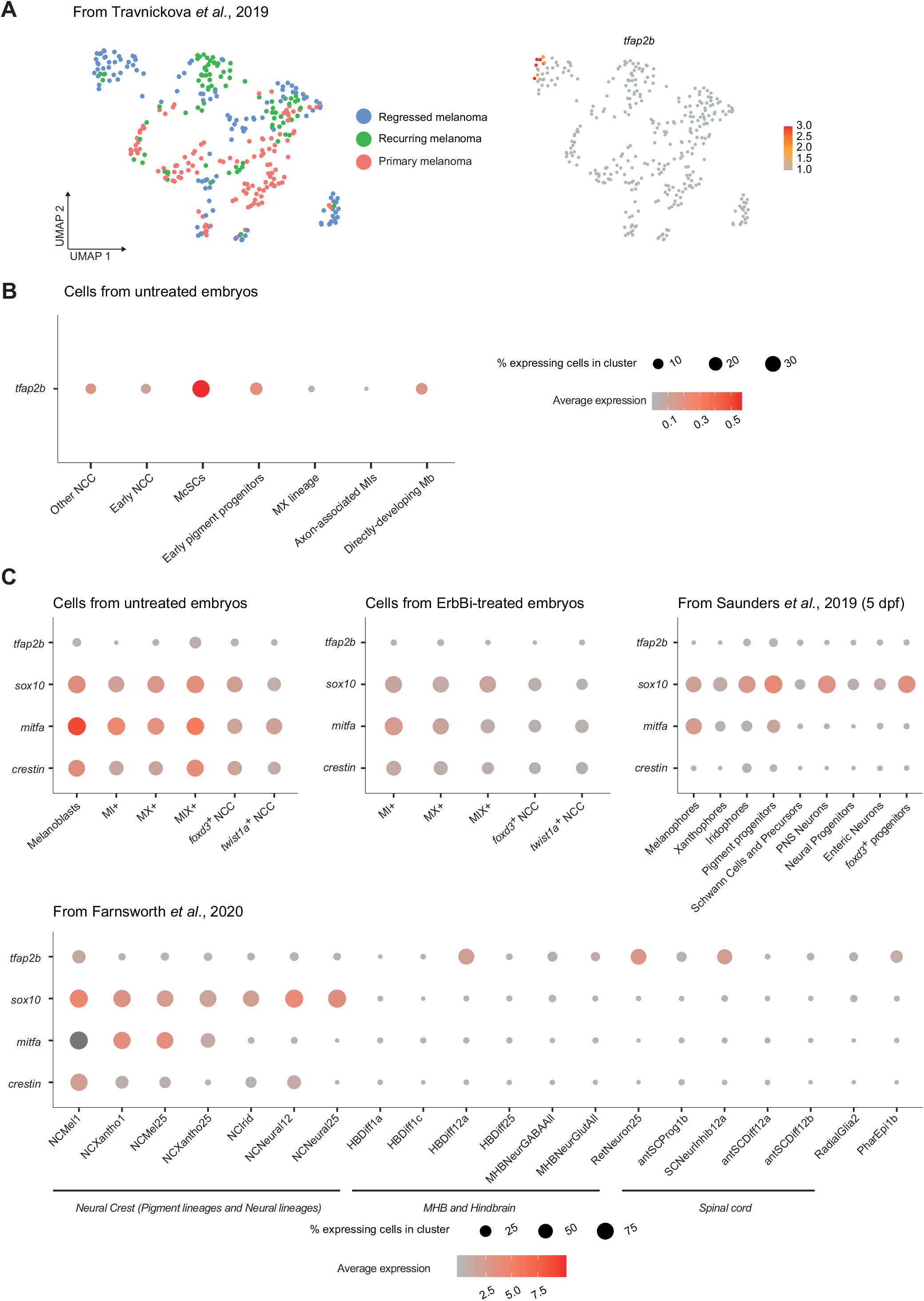
*tfap2b* expression in development and melanoma residual disease. **A.** *tfap2b* expression in a cluster of cells from melanoma residual disease. Left: UMAP of cells retrieved from GSE136900 (Travnickova et al., 2019) pseudocolored with cell-origin. Right: The same UMAP with color change based on log_2_ mRNA expression of *tfap2b*. *tfap2b* is expressed only in a subpopulation of cells from residual disease in the regressed melanomas. **B.** Dot-plot representing the average expression of *tfap2b* across lineages identified by integrated pseudotime analysis (Figure 3C). **C.** Dot-plots representing the average expression of *tfap2b*, *mitfa*, *sox10* and *crestin* in untreated and ErbBi-treated embryo datasets, and in published datasets (Farnsworth et al., 2020; Saunders et al., 2019).

**Figure S6.**
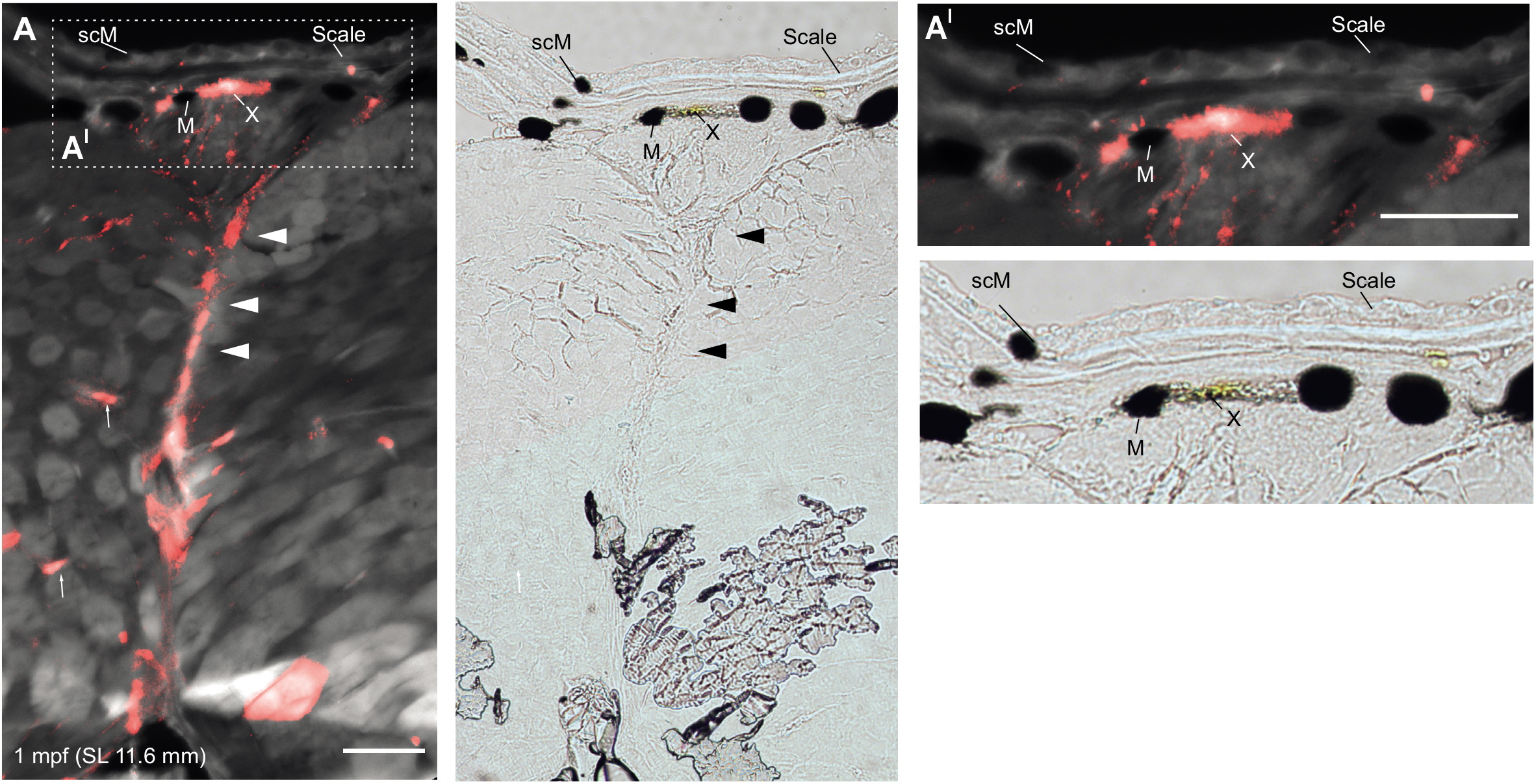
Cryo-sections of juvenile zebrafish reveal *tfap2b+* progenitors in the skin and along nerves. Fate mapping *tfap2b+* progeny in cross sections of juveniles by cryo-section (from Figure 7). mCherry+ cells migrate along nerves (arrow heads) to form skin associated clones in the dorsal stripe (as labelled). mCherry+ precursors can be seen as associated with (possibly) minor axonal projections lateral to the main nerve (small arrows). Melanocytes (M) and xanthophores (X) are identifiable in bright field by the black and yellow pigment granules. The dashed box indicates the region magnified at right. Left panel: merged view (mCherry, red; GFP, gray); right panel: brightfield. SL: Standard Length. scM: scale melanocyte. Scale bar: 50 μm. Section thickness: 40μm.

### Supplementary tables

**Table S1:** Metrics, clustering information, and cell states for all cells.

**Table S2:** Cluster markers. Comparison between indicated cluster relative to pooled cells of all other clusters.

**Table S3:** Differential expression analysis of McSCs vs pigment lineage cells.

**Table S4:** Pathway analysis of DE genes from McSCs vs pigment lineage cells.

**Table S5:** Differential expression analysis of McSCs vs Early pigment progenitors.

**Table S6:** List of Tfap2b targets.

**Table S7:** Differential expression analysis of untreated versus ErbBi-treated MIX+ cells.

